# Myelin density dictates region-specific vulnerability of oligodendrocyte lineage cells during aging

**DOI:** 10.64898/2025.12.02.691836

**Authors:** Aladdin Skaf, Jaime Eugenin von Bernhardi, Leda Dimou

## Abstract

Aging of the central nervous system (CNS) leads to a progressive decline in numerous physiological functions. This decline can be attributed in part to alterations within the oligodendrocyte lineage, which comprises myelinating oligodendrocytes (OLs) and their progenitors, NG2-glia, that play a central role in maintaining homeostasis and ensuring proper myelin turnover. While NG2-glia are distributed throughout the CNS, OLs are enriched in highly myelinated regions, implying spatially heterogeneous requirements for NG2-glia proliferation and differentiation. Consequently, age-related impairments in these progenitor functions may differentially compromise oligodendrogenesis and myelin maintenance across distinct CNS compartments. Yet, the spatial and temporal dynamics of aging-associated alterations within the oligodendrocyte lineage remains insufficiently characterized.

To address this gap, we investigated the effects of aging across three age groups in two anatomically adjacent but functionally distinct CNS regions, the cortical gray matter (GM) and the corpus callosum (CC).

We demonstrated that aging is accompanied by a marked decline in the NG2-glia population. Aging NG2-glia displayed cell cycle dysregulation, characterized by reduced proliferative and differentiative capacity and accompanied by increased expression of cyclin-dependent kinase inhibitors (CDKIs), indicating disrupted homeostatic regulation. These alterations were most pronounced in highly myelinated regions, which also exhibited a stronger shift toward an age-associated inflammatory milieu. In parallel, we observed substantial accumulation of myelin debris and impaired phagocytic clearance in these myelin-dense areas. Moreover, we identified a selective loss of myelinating OLs in the CC, a phenomenon not detected in the gray matter (GM).

Together, our findings delineate a tight interdependence between regional myelin density, inflammatory status, and the vulnerability of oligodendrocyte lineage cells to aging. These highlight multiple entry points of potential therapeutic intervention to mitigate CNS aging.

## 1. Introduction

Aging is a gradual physiological process that impairs cellular and tissue function, reducing the capacity of an organism to respond and adapt to metabolic and environmental stress. It is characterized by well-established hallmarks, such as cellular senescence and stem cell exhaustion (López-Otín et al., 2013, 2023).

In the central nervous system (CNS), aging impairs multiple brain functions and is associated with a progressive decline in cognitive, motor and adaptive abilities, like (motor) learning (Bishop et al., 2010; Inoue & Nishimune, 2023). While research has predominantly focused on the neuronal aspect of CNS aging, glial cells, which constitute roughly half of the cells of the CNS, play equally critical roles, yet remain comparatively understudied (Allen & Lyons, 2018). For instance, the age-dependent effects on neurogenesis – a process that occurs during lifetime in the adult neurogenic niches such as the dentate gyrus (DG) and the subependymal zone (SEZ) – were well studied (Apple et al., 2017). In contrast, the impact of aging on glia, particularly oligodendrocyte lineage cells and oligodendrogenesis, remains poorly understood.

Oligodendrocyte lineage cells comprise NG2-glia, also known as oligodendrocyte progenitor cells (OPCs) and oligodendrocytes (OLs), the myelinating cells of the CNS (Viganò & Dimou, 2016a). NG2-glia are widely and homogenously distributed across the brain, proliferate throughout life, and serve as the sole progenitor pool outside the neurogenic niches (Dimou et al., 2008a; Young et al., 2013). The main function of NG2-glia is to generate OLs via differentiation, a process termed oligodendrogenesis, which occurs lifelong while maintaining strict homeostatic control over their density (Hill et al., 2018; Hughes et al., 2013; Viganò & Dimou, 2016a). In contrast, OL distribution is regionally heterogeneous across the CNS (Valério-Gomes et al., 2018), with higher densities in myelin-rich areas, such as the corpus callosum (CC), where oligodendrogenesis is elevated to meet local myelination demands (Hardy & Reynolds, 1991; Kuhn et al., 2019; Viganò & Dimou, 2016b). Under physiological conditions, oligodendrogenesis supports continuous myelination of new axonal segments, allowing lifelong remodeling and adaptation of existing myelin patterns in an experience-dependent manner (Dimou & Götz, 2014; Gibson et al., 2014; Hughes et al., 2018). Moreover, experimental modulation of oligodendrogenesis has been shown to enhance multiple brain functions, including cognition and learning (Bacmeister et al., 2020; Eugenin von Bernhardi & Dimou, 2022).

During aging, NG2-glia exhibit reduced proliferation and differentiation rates, resulting in fewer newly generated OLs (Sim et al., 2002; Young et al., 2013; Zhu et al., 2011). Furthermore, newly formed OLs produce shorter and fewer myelin sheaths (Lasiene et al., 2009) while myelin degradation and accumulation of myelin debris within microglia increase over time in the superficial cortical areas as demonstrated by *in vivo* imaging (Hill et al., 2018). Critically, these processes are not uniform across the CNS; some regions are more susceptible to age-associated decline (Raz et al., 2005; Zhou et al., 2023), highlighting the need to investigate the regional and temporal dynamics of aging in oligodendrocyte lineage cells.

Here, we investigated the effects of aging on NG2-glia proliferation, oligodendrogenesis, and myelination in two adjacent yet distinct brain regions, the cortical gray matter (GM) of the motor cortex and the underlying white matter tract of the CC. Our results revealed region specific differences in NG2-glia sensitivity to aging. In the GM, homeostatic disruption was largely confined to deep layers, whereas the CC exhibited an earlier and more pronounced loss in NG2-glia density. Moreover, we showed that the decrease in the proliferative capacity of NG2-glia occurs gradually in the GM with age, while this decline was abrupt in the CC and occurred earlier in life. Correspondingly, oligodendrogenesis decreased primarily in myelin-dense regions, like the deep layers of the GM and the CC, whereas intermediate and superficial GM layers remained largely unaffected.

Aging also induced a pronounced inflammatory shift in the CC, characterized by microgliosis and infiltration of peripheral immune cells, coinciding with a selective loss of mature, myelinating OLs (MOLs). In contrast, in the GM, we found a consistent increase in MOLs throughout lifespan and no changes in the density of microglia. Across both regions, microglia displayed age dependent morphological alterations, including cytoplasmatic enlargements in the shape of pockets often containing myelin inclusions, indicative of ongoing engulfment. Notably, the age-dependent morphological changes of microglia preceded myelin debris inclusions, and engulfment events increased progressively with age, reflecting impaired myelin debris clearance.

Together, our findings reveal a robust interplay between myelin load, inflammatory state, and oligodendrocyte lineage cells vulnerability during brain aging. Our results highlight how region- and layer-specific differences in oligodendrogenesis and microglial activity contribute to age-associated white matter decline, providing a framework for understanding the cellular mechanisms underlying CNS aging.

## 2. Materials and methods

### 2.1 Animals

NG2CreER^T2^xCAG-eGFP (Huang et al., 2014), Plp-dsRed (Hirrlinger et al., 2005) and GPR17-iCreER^T2^xCAG-eGFP transgenic (Viganò et al., 2016) mouse lines were utilized for the young (12-16 weeks), middle-aged (50-54 weeks), and aged (80-114 weeks) groups, respectively. Animals were kept in a 12/12 h dark-light cycle between 20 and 24°C. Genomic PCRs confirmed the genotype of animals, as described in previous publications. Both sexes were used.

All conducted experiments followed the guidelines on using animals and humans in neuroscience research, revised and approved by the Society of Neuroscience and licensed by the State of Upper Bavaria and Baden Wuerttemberg.

### 2.2 Tamoxifen induction and EdU administration

Tamoxifen was dissolved in ethanol and subsequently diluted with corn oil (10% ethanol, 90% corn oil) to a final concentration of 40 mg/mL. Tamoxifen (Sigma-Aldrich #T5648-5G) was administered to the NG2CreER^T2^xCAG-eGFP line three times every other day by oral gavage at a dose of 10 mg per 30 g of body weight.

For the proliferation experiments, adult NG2CreER^T2^xCAG-eGFP and GPR17-iCreER^T2^xCAG-eGFP mice received 5-ethynyl-2′-deoxyuridine (EdU) (Thermo Fisher #C10340) in their drinking water at a concentration of 0.2 mg/mL supplied with 1% sucrose for two consecutive weeks.

### 2.3 Histology and immunofluorescence

For the NG2CreER^T2^xCAG-eGFP cohort, animals were anesthetized with an i.p. injection of a drug combination containing ketamine (100 mg/kg) and xylazine (10 mg/kg) diluted in a 0.9% NaCl solution. After anesthesia, transcardial perfusion was performed, pumping 25 ml of PBS and 50 ml of 4% PFA in PBS as a fixative subsequently. Later, brains were extracted and immersed in 4% PFA for 2 h for postfixation. Next, brains were immersed in 30% sucrose in PBS until sinkage at 4 °C for cryoprotection. For the Plp-dsRed cohort, animals were sacrificed using the cervical dislocation. Next, brains were dissected and postfixed with 4%PFA for 6 hours. Later, brains were transferred into 30% saccharose for cryoprotection. Free-floating 30μm thick sections were prepared and kept in PBS at 4°C. For staining, slices were blocked and permeabilized in PBS 0.5% Triton-X™ and 2% bovine serum albumin (BSA) for 1 h. After washing with PBS, sections were incubated in specific combinations of the following primary antibodies diluted in block/perm solution ON at 4 °C:

Rabbit αNG2 (1:500, Merck #AB5320), mouse IgG2b αCC1 (1:100, Merck #OP80), mouse αGST-π (1:250, BD Pharmingen #610719), chicken αGFP (1:500, Aves #GFP-1010), rat αAN2 (1:200, kindly provided by Prof. Jacky Trotter, University of Mainz), rat αPDGFRα (1:200, Thermo Fisher #12-1401-81), guinea pig αIba1 (1:500, Synaptic Systems #234308), mouse αRFP (1:500, Thermo Fischer #MA5-15257), rat αp19ARF/CDKN2A (1:250, Novus Biologicals #NB200-169SS), mouse αp16 CDKN2A (1:250, Santa Cruz #sc-1661), mouse αMAG (1:1000, Merck #MAB1567), rabbit αKi67 (1:250, Thermo Fisher #MA514520), rabbit αLAMP1 (1:500, Abcam #ab208946), rabbit αNAV1.6 (1:500, Alomone Labs #ASC-009), and mouse IgG1 αCaspr (1:500, NeuroMab #1308-1381).

For p16 CDKN2A, p19 ARF/CDKN2A and Ki67 detection, slices underwent an antigen retrieval step with a heated citrate buffer. A 1,5 ml Eppendorf filled with 1 ml citrate buffer was pre-heated on a thermocycler to 96 °C. Samples were added to the prewarmed citrate, put back into the thermocycler for 10 minutes, and then placed at RT for 10 minutes to cool down. Slices were then washed three times with 1xPBS and proceeded with incubation with the primary antibodies, as mentioned above.

The next day, slices were washed three times in PBS and incubated in a combination of the following secondary antibodies diluted in block/perm solution for 2 h at room temperature with shaking: αChicken Alexa Fluor® 488 (1:250, Invitrogen #A-11039), αMouse IgG2b Alexa Fluor® 647 (1:250, Invitrogen #A-21242), αRabbit Alexa Fluor® 488 (1:250), αMouse IgG Cy® 3 (1:250, Dianova #115-165-166), αRabbit Cy® 3 (1:250, Dianova #711-165-152), αMouse Alexa Fluor 568 (1:250, Thermo Fisher #A11004), αGuinea pig IgG Cy® 3(1:250, Dianova #706-165-148), αMouse IgG2b Alexa Fluor® 555 and αRat Cy® 3 (1:250, Dianova #112-165-167), αRabbit.Cy® 5 (1:250, Dianova #711-175-152).

For EdU detection, all other immunostainings were performed before, and slices were treated with the highly specific click reaction kit (Click-iT™ Alexa Fluor® 647, Thermo Fisher Scientific #C10340) as indicated by the manufacturer. Finally, slices were counterstained with DAPI (1:1000 in PBS) and mounted on a slide with aqua-poly/mount (Polysciences #18606-5).

### 2.4 Terminal deoxynucleotidyl transferase-dUTP nick end labeling

For the terminal deoxynucleotidyl transferase–dUTP nick end labeling (TUNEL) assay, we used the Click-iT™ Plus TUNEL Assay kit (Invitrogen, #C10619) following the manufacturer’s instructions. Briefly, free-floating brain sections were fixed in 4% PFA for 15 min at 37 °C, washed in PBS, and permeabilized with Proteinase K for 15 min. Sections were then washed in PBS and rinsed in deionized water.

To initiate DNA end labeling, 100 µL of TdT Reaction Buffer (Component A) was added to each section, ensuring complete coverage, and incubated for 10 min at 37 °C. The TdT reaction mixture was subsequently prepared according to the manufacturer’s protocol using TdT Reaction Buffer (Component A), EdUTP (Component B), and TdT enzyme (Component C). Sections were transferred into the TdT reaction mix and incubated for 60 min at 37 °C.

After incubation, sections were rinsed in deionized water and washed with 3% BSA/0.1% Triton™ X-100 in PBS. The Click-iT™ Plus detection reaction was then prepared using Click-iT™ Plus TUNEL Supermix and 10× Reaction Buffer Additive.

Sections were immersed in 50 µL of the Click-iT™ Plus reaction cocktail and incubated for 30 min at 37 °C protected from light. Following a final wash in PBS, sections were processed for subsequent immunostaining.

### 2.5 Image acquisition and analysis

Images were taken with a Leica SPE confocal microscope with a 40x magnification, manually processed, and analyzed in Fiji (ImageJ) and Leica’s LAS X Life Science. Only cells containing a visible DAPI signal were considered.

For the analysis of the cortical GM, confocal images of a whole column of the sensorimotor cortex containing all cortical layers were taken and three columns per animal were analyzed. Each column was further divided with ImageJ into three equal areas (bins) and subsequently numbered from superficial to deep as bin-1, -2, and -3.

Nine pictures of different areas were taken and analyzed for the CC analysis. For both acquisitions, at least three slices per animal were quantified.

To measure nodal and paranodal compartments, confocal imaging was performed in three matched callosal regions per animal. The midline was used as a fixed anatomical landmark, and three consecutive lateral fields of view were acquired to ensure consistent anatomical sampling across animals. Within each image, 50 nodal and paranodal segments were randomly sampled and measured, yielding 150 measurements per animal.

All quantifications were done manually with a cell counter plugin in Fiji (ImageJ). Additionally, the 3D viewer plugin was used for better visualization of pocketed microglia morphology and their contents.

The Keyence All-in-one Fluorescence Microscope (BZ-X810) was used to quantify TUNEL-positive cells. Three hemispheres per animal were analyzed, and TUNEL-positive cells were quantified manually using multi-channel visualization. The areas examined were the cortical GM and the CC.

All analyses were conducted in a blinded manner.

### 2.6 Statistics

All quantitative data are reported as mean ± SEM. No statistical method was used to predetermine sample size. Instead, sample sizes were based on previous studies in the field. The Shapiro–Wilk test was used to assess the normality of each dataset. No exclusion criteria were applied.

To compare means among three independent groups, either a one-way unpaired analysis of variance (ANOVA) with Tukey’s multiple comparison post hoc test (for parametric data) or the Kruskal–Wallis test with Dunn’s multiple comparison (for non-parametric data) was used. For grouped analyses—where three independent age groups were further subdivided by a second independent variable—two-way unpaired ANOVA with Tukey’s multiple comparison post hoc test (parametric) was performed. All statistical tests were conducted using a two-tailed manner.

For correlation analyses between age and various glial cell densities, either a simple linear regression with Pearson’s correlation coefficient (for parametric data) or Spearman’s rank correlation (for non-parametric data) was applied, depending on data distribution. In cases where the data exhibited a non-linear trend, a log-normal regression model was employed to account for the distribution and improve the goodness of fit. Data were considered significant at *p*-value ≤ 0.05 = *, *p*-value ≤ 0.01 = **, and *p*-value ≤ 0.001 = ***. All statistical tests were performed using GraphPad Prism 10.

## 3. Results

### Aging disrupts the homeostatic control of NG2-glia

NG2-glia are characterized by their widespread, homogenous distribution and strict homeostatic regulation of their population density across the brain (Hughes et al., 2013). To assess how aging affects this regulation, we quantified NG2-glia numbers in the GM of the cerebral cortex and the adjacent CC. Our results reveal that aging coincides with a subtle yet significant reduction in NG2-glia density in the GM (young vs. aged *p=0.0016*; Figure 1 A). Although the middle-aged group did not differ significantly from either young or aged animals, the NG2-glia numbers suggest a gradual age-dependent decline of NG2-glia density in the GM (Figure 1 A).

**Figure 1:**
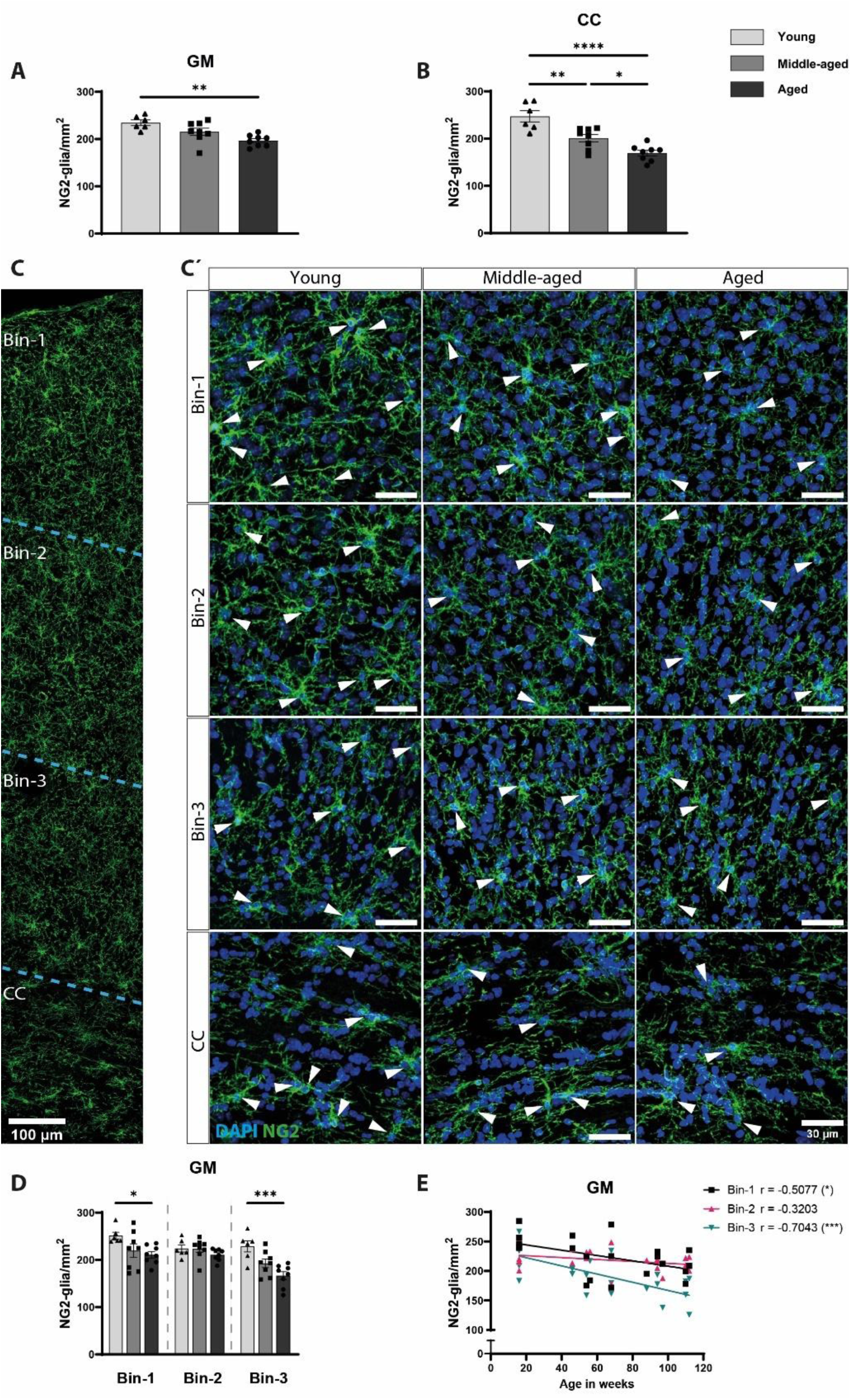
The age-dependent alterations of NG2-glia density. (A) Numbers of NG2-glia in the gray matter of the motor cortex (GM) and (B) the adjacent corpus callosum (CC). (C) Immunostaining image from a column taken across the motor cortex showcasing the division of the GM into three equal bins and the adjacent CC. The scale bar represents 100µm. (C’) Immunostaining with DAPI (blue) and NG2 (green) from bins-1, 2 and 3 and CC of young, middle-aged and aged mice showing NG2-glia (white arrowheads). The scale bar represents 30µm. (D) The distribution of NG2-glia in bins 1-3 of the GM in young, middle-aged and aged groups (n=6, n=8 and n=8, respectively). All data are plotted as mean ± SEM. Each dot represents an individual animal. Statistical differences were determined by unpaired one-way ANOVA with Tukey’s post-hoc multiple comparisons tests. *p* value ≤ 0.05 = *, *p* value ≤ 0.01 = **, *p* value ≤ 0.001 = *** and *p* value ≤ 0.0001 = ****. (E) Linear correlation analysis between the age and the density of NG2-glia in the bins of the GM. n=22 (n=6, n=8 and n=8 young, middle-aged, and aged respectively). A linear correlation analyzed data and fit to a linear regression for each bin.

In the CC however, NG2-glia loss was more pronounced. Density was significantly reduced not only in the aged group compared to the young group (young vs. aged *p<0.0001*; Figure 1 B) but was already diminished in the middle age group (young vs. middle-aged *p=0.0036*; Figure 1 B), with a further decline in aged animals (middle-aged vs. aged *p=0.0293*; Figure 1 B). Hence, a more pronounced loss of NG2-glia was observed in the aging CC.

Consistent with these observations, linear correlation analysis revealed a significant negative correlation between age and NG2-glia density in both regions (GM: r= - 0.7184, *p=0.0002*; CC: r= -0.7697, *p<0.0001*; Suppl. Figure 1 A-B).

To determine whether specific cortical layer depths exhibit differential vulnerability, we analyzed the distribution of NG2-glia across three equal-sized bins of the GM, bin-1 being the superficial, outermost and bin-3 being the deepest, innermost (Figure 1 C). Remarkably, aging affected NG2-gia density in a layer dependent manner. In superficial (bin-1) and deep layers (bin-3), NG2-glia numbers declined significantly (bin-1: young vs. middle-aged *p=0.0297*; bin-3: young vs. aged *p=0.002;* Figure 1 D). In contrast, intermediate layers of the GM (bin-2) showed no significant age-related changes in the density of NG2-glia (Figure 1 D). Correspondingly, the strongest negative linear correlation between age and NG2-glia density was observed in the deepest layers of the GM, followed by the superficial layers of the GM (bin-1: r= - 0.5077 *p=0.0159*; bin-3: r= -0.7043 *p=0.0003*; Figure 1 E). Together, these findings demonstrate that aging disrupts NG2-glia homeostasis in both the GM and CC, but with distinct temporal and spatial patterns. This is shown by the CC exhibiting an earlier and more pronounced decline of NG2-glia, highlighting its heightened vulnerability compared to the GM of the motor cortex.

### The regenerative capacity of NG2-glia declines with age

To better understand the dynamics behind the age dependent loss of NG2-glia, we next examined how aging alters their proliferative capacity We first quantified the number of actively proliferating NG2-glia (PDGFRα^+^Ki67^+^). In the GM, proliferation was markedly reduced in the aged group compared with both young and middle-aged groups, whereas no significant difference was observed between young and middle-aged mice (young vs. aged *p=0.0159*, middle-aged vs. aged *p=0.0173*; Suppl. Figure 1 C-D). In the CC however, the decline in active proliferation occurred earlier and more abruptly: NG2-gia proliferation was already significantly reduced in the middle-aged group and continued to drop further in the aged group (young vs. middle-aged *p=0.0018*, middle-aged vs. aged *p=0.0409*, young vs. aged *p<0.0001*; Suppl. Figure 1 E). Furthermore, correlation analysis further revealed a strong negative correlation between age and proliferation of NG2-glia in both regions (GM: r=-0.7997, CC: r=-0.7852; Suppl. Figure 1 F-G), further mirroring the accelerated vulnerability of NG2-glia in the CC described before.

To further assess proliferative dynamics, we traced cycling NG2-glia using an EdU pulse-retention paradigm (Figure 2 A-B). EdU allows the tracing of proliferated cells by integrating into the DNA during the cell cycle. In the GM, the number of EdU^+^ NG2-glia (PDGFRα^+^EdU^+^) declined significantly with age (young vs. aged *p=0.0257*; Figure 2 C). Layer-specific analysis revealed a significant negative correlation across all GM bins, with the strongest association in superficial areas (bin-1: r=-0.7228 *p=0.0005*; bin-2: r=-0.6326 *p=0.0037*; bin-3: r=-0.5095 *p=0.0258*; Figure 2 D). In the CC, EdU^+^ NG2-glia numbers also declined significantly with age, but consistent with Ki67 data, this reduction was already evident in middle-aged animals and remained low thereafter (young vs. middle-aged *p<0.0001*, young vs. aged *p<0.0001*; Figure 2 E). Correlation analysis confirmed a significant age-dependent decline in the numbers of proliferated NG2-glia (PDGFRα^+^EdU^+^) (GM: r= -0.7165, CC: r= -0.6006; Suppl. Figure 1 H-I).

**Figure 2:**
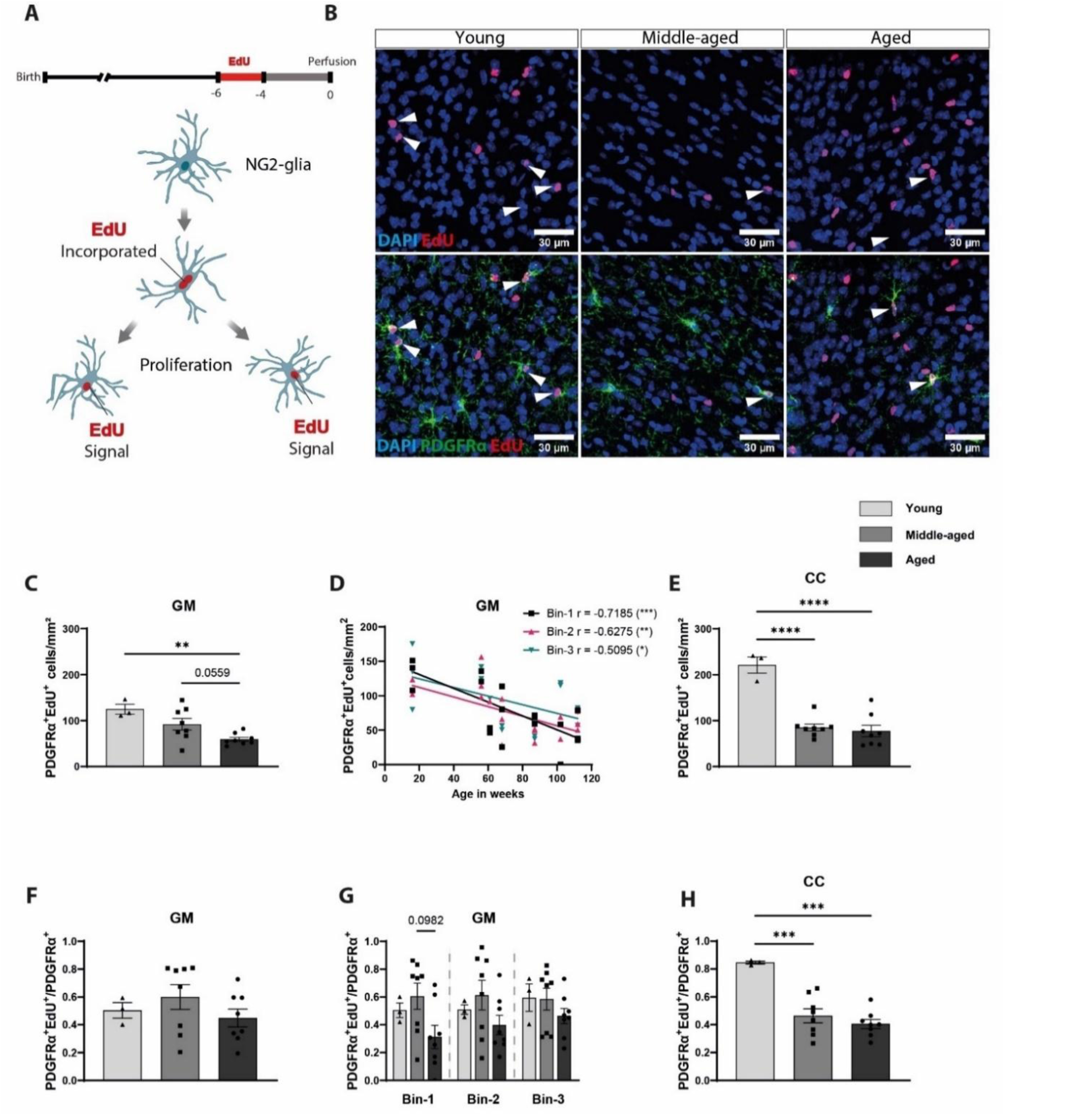
The proliferative capacity of NG2-glia declines with age. (A) A scheme showing the EdU pulse in red (2 weeks) and the retention period in gray (4 weeks) for tracing proliferated NG2-glia. (B) Immunostaining images of proliferated cells in red (EdU^+^) and NG2-glia in green (PDGFRα^+^), arrowheads show proliferated NG2-glia (PDGFRα^+^EdU^+^). The scale bar represents 30µm. (C) Numbers of proliferated NG2-glia in the cortical GM. (D) Simple linear regression with Pearson’s correlation analysis between age and numbers of proliferated NG2-glia (PDGFRα^+^EdU^+^) in different bins of the GM. (E) Quantification of proliferated (dark blue sub-column) and non-proliferated NG2-glia (light blue sub-column) in the CC. (F) The fraction of proliferated NG2-glia in the cortical GM, (G) across different layers of the GM, and (H) in the CC. All data are plotted as mean ± SEM. Each dot represents an individual animal. Two-way ANOVA determined statistical differences with Tukey’s post-hoc multiple comparisons tests for graphs (E and G) and one-way ANOVA with Tukey’s post-hoc multiple comparisons tests for graphs (B-C). *p* value ≤ 0.05 = *, *p* value ≤ 0.01 = **, *p* value ≤ 0.001 = *** and *p* value ≤ 0.0001 = ****.

Knowing that NG2-glia density itself declines with age, we normalized the numbers of cycling NG2-glia to the total NG2-glia pool. Interestingly, this revealed that the fraction of cycling NG2-glia (PDGFRα^+^EdU^+^/ PDGFRα^+^) remained stable across age groups in the GM, including across individual cortical bins (Figure 2 F-G). In contrast, the fraction of cycling NG2-glia in the CC dropped significantly with age, namely from 84.61% in young, animals to 46.35% in middle-aged, stabilizing at 40.51% in aged mice (young vs. aged *p<0.0001*, young vs. middle-aged *p=0.0003*; Figure 2 H). These data suggest that CC-resident NG2-glia undergo substantive age-associated alterations in cell cycle entry of progression that exceed those observed in the GM.

Given the known regional differences in NG2-glia cell cycle length (Dimou et al., 2008a) and the central role of cell cycle halt during aging (López-Otín et al., 2023), we examined the expression of the cell cycle regulators p16, p19, or both, in genetically labeled NG2-glia (NG2CreER^T2^xCAG-eGFP mice (Huang et al., 2014)). The proportion of recombined NG2-glia expressing the cell cycle regulators p16, p19 or both (GFP^+^p16^+^p19^+^) increased significantly with age in both the GM and the CC (GM: young vs. aged *p=0.*0122, middle-aged vs. aged *p=0125*; CC: middle-aged vs. aged *p=0.0241*; Suppl. Figure 2 A-C). A similar pattern emerged when normalizing p16 and p19-expressing cells to the total recombined population (GFP^+^p16^+^p19^+^/GFP^+^) (GM: young vs. aged *p=0.0542*, middle-aged vs. aged *p=0.0159*; CC: middle-aged vs. aged *p=0.0102*; Suppl. Figure 2 D-E). These results indicate that aging progressively impairs the regenerative capacity of NG2-glia by reducing both proliferation and cell cycle progression, with a markedly earlier and more pronounced decline in the myelin-rich CC. This region-specific disruption of NG2-glia homeostasis likely contributed to the heightened vulnerability of white matter during aging.

### Oligodendrogenesis is subjected to aging in a spatially dependent manner

NG2-glia can terminally differentiate into oligodendrocytes (OLs) and retain this capacity throughout life (Dimou et al., 2008). However, the production of newly generated OLs (NGOLs) declines with age (Sim et al., 2002). Whether this reduction occurs uniformly across the brain, remains unresolved. To dissect potential spatial heterogeneity, we performed a fate-mapping experiment of the progeny of NG2-glia in the inducible NG2CreER² x GFP mouse line (Huang et al., 2014). After induction with tamoxifen, a two-week EdU pulse was administered *ad libitum,* followed by a four-week retention period. This experimental strategy enables the identification of two modes of oligodendrogenesis, namely the direct (GFP^+^CC1^+^EdU^−^) and the indirect, proliferation-dependent differentiation (GFP^+^CC1^+^EdU^+^/GFP^−^CC1^+^EdU^+^) (Figure 3 A).

**Figure 3:**
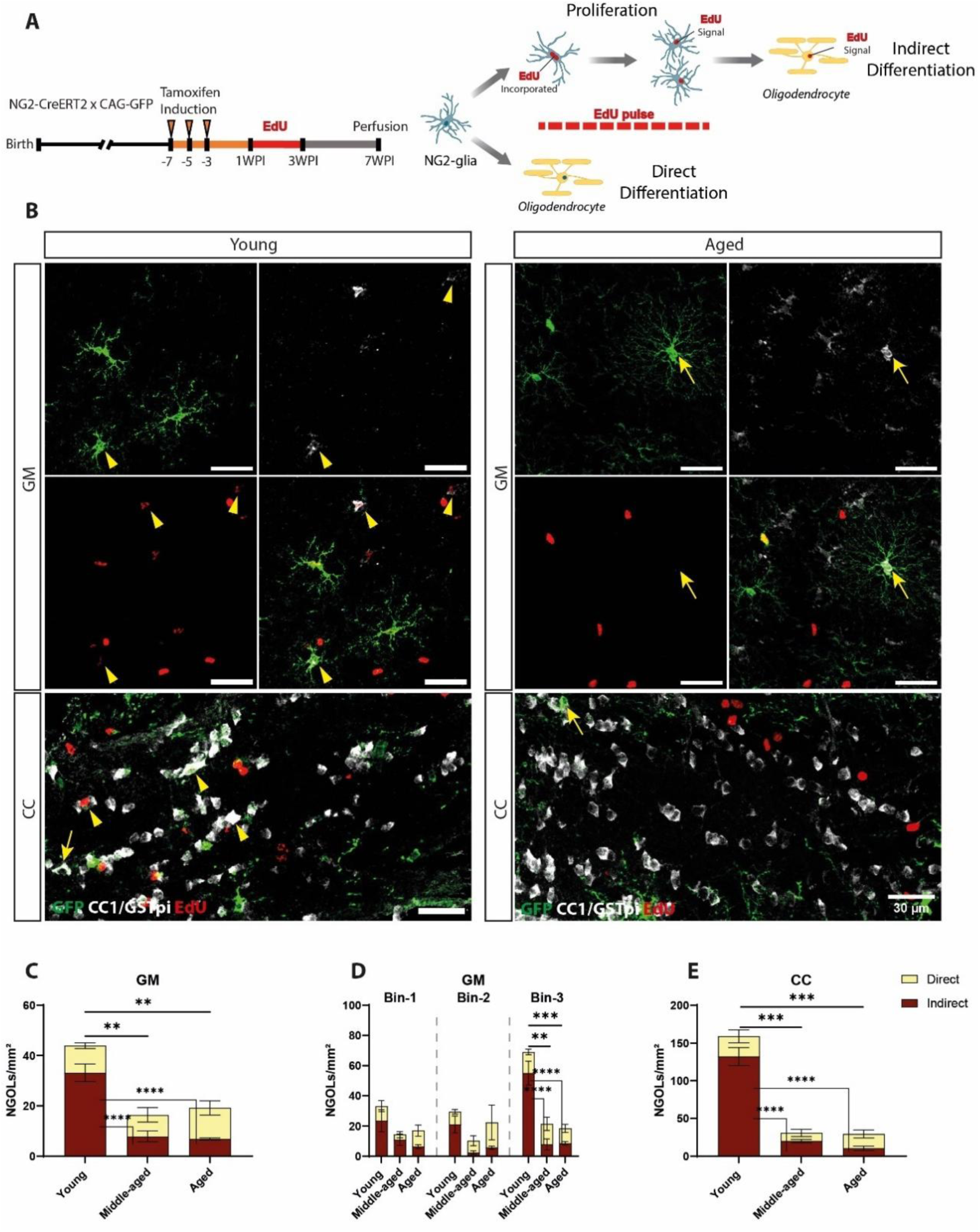
Age-related alterations in oligodendrogenesis are spatially restricted to areas of high myelin demand. (A) Schematic overview of the fate-mapping experiment of the progeny of NG2-glia in the NG2-CreERT^2^ x GFP mouse line; Left: the timeline of the experiment with the induction of the Cre-recombinase with Tamoxifen shown in orange, the administration of EdU pulse is highlighted in red (2 weeks) followed by retention period shown in gray (4 weeks). Right: scheme showing the direct (EdU^−^) and the indirect (EdU^+^) differentiation modalities of NG2-glia. (B) Immunohistochemical staining of recombined cells (GFP), oligodendrocytes (CC1/GSTpi) and proliferated cells (EdU) from young and aged samples. Arrowheads show indirectly formed NGOLs (GFP^+/-^CC1^+^EdU^+^). Arrows show directly formed NGOLs (GFP^+^CC1^+^EdU^−^). The scale bar represents 30µm. (C) Quantification of directly (dark blue sub-column) and indirectly NGOLs (light blue sub-column) in the cortical GM. (D) Quantification of directly (dark blue sub-column) and indirectly NGOLs (light blue sub-column) in different bins of the GM. (E) Quantification of directly (dark blue sub-column) and indirectly NGOLs (light blue sub-column) in the CC. All data are plotted as mean ± SEM. n=3 in each age group. Two-way ANOVA determined statistical differences with Tukey’s post-hoc multiple comparisons tests *p* value ≤ 0.01 = **, *p*-value ≤ 0.001 = *** and *p*-value ≤ 0.0001 = ****.

In the GM, aging selectively impaired the indirect modality of oligodendrogenesis, where a proliferative step precedes differentiation (indirect: young vs. middle-aged *p<0.0001,* young vs. aged *p<0.0001*; Figure 3 C). In contrast, direct differentiation remained stable across age groups (Figure 3 C). Interestingly, layer specific analysis revealed that this decline was confined for the deep cortical layers of the GM (bin-3), whereas superficial layers showed no significant age-related reduction (indirect: bin-3 young vs. middle-aged *p<0.0001* and young vs. aged *p<0.0001*; Figure 3 D).

A similar pattern emerged in the CC: the number of indirectly NGOLs was profoundly reduced in both middle-aged and aged animals, whereas direct NGOLs numbers remained unchanged (indirect: young vs. middle-aged *p<0.0001* and young vs. aged *p<0.0001*; Figure 3 F). These findings indicate that the age-related impairment is particularly pronounced in regions with high density of OLs, namely the deep layers of the GM and the CC, mirroring the region-specific decline in NG2-glia proliferation described above.

We next analyzed the numbers of newly generated myelinating OLs (NGMOLs) in the GM (GFP^+^MAG^+^ and GFP^+/-^MAG^+^EdU^+^). NGMOL density was significantly reduced in middle-aged and aged animals compared to young mice, with no further decline between middle-aged and aged groups (young vs. middle-aged *p=0.0144*, young vs. aged *p=0.0329*; Suppl. Figure 3 A-C). An equivalent quantification in the CC was not feasible due to dense MAG labeling within this area. Together, these data demonstrate that although oligodendrogenesis declines with age, particularly in myelin-rich regions, a subset of NGOLs retains the ability to integrate into the myelination network throughout life.

### Loss of myelinating OLs in the aging corpus callosum

Given that oligodendrogenesis continuously supplies NGOLs into the myelination network (Hill et al., 2018; Rivers et al., 2008; Young et al., 2013), we next examined how the age-related decline in NG2-glia proliferation and differentiation affects the density of myelinating OLs (MOLs). We used the Plp-dsRed reporter mouse line (Hirrlinger et al., 2005), we quantified MOL density in both the GM and the adjacent CC.

In the GM, MOL numbers increased steadily across the lifespan, indicating sustained incorporation of new OLs despite reduced proliferative activity of NG2-glia. MOL density rose significantly from young to middle aged and further into old age (young vs. middle-aged *p<0.0001*; middle-aged vs. aged *p=0.0034*; young vs. aged *p<0.0001*; Figure 4 A-B). This was supported by a prominent positive correlation between age and MOL abundance (r=0.8610 *p<0.0001*; Suppl. Figure 1 D).

**Figure 4:**
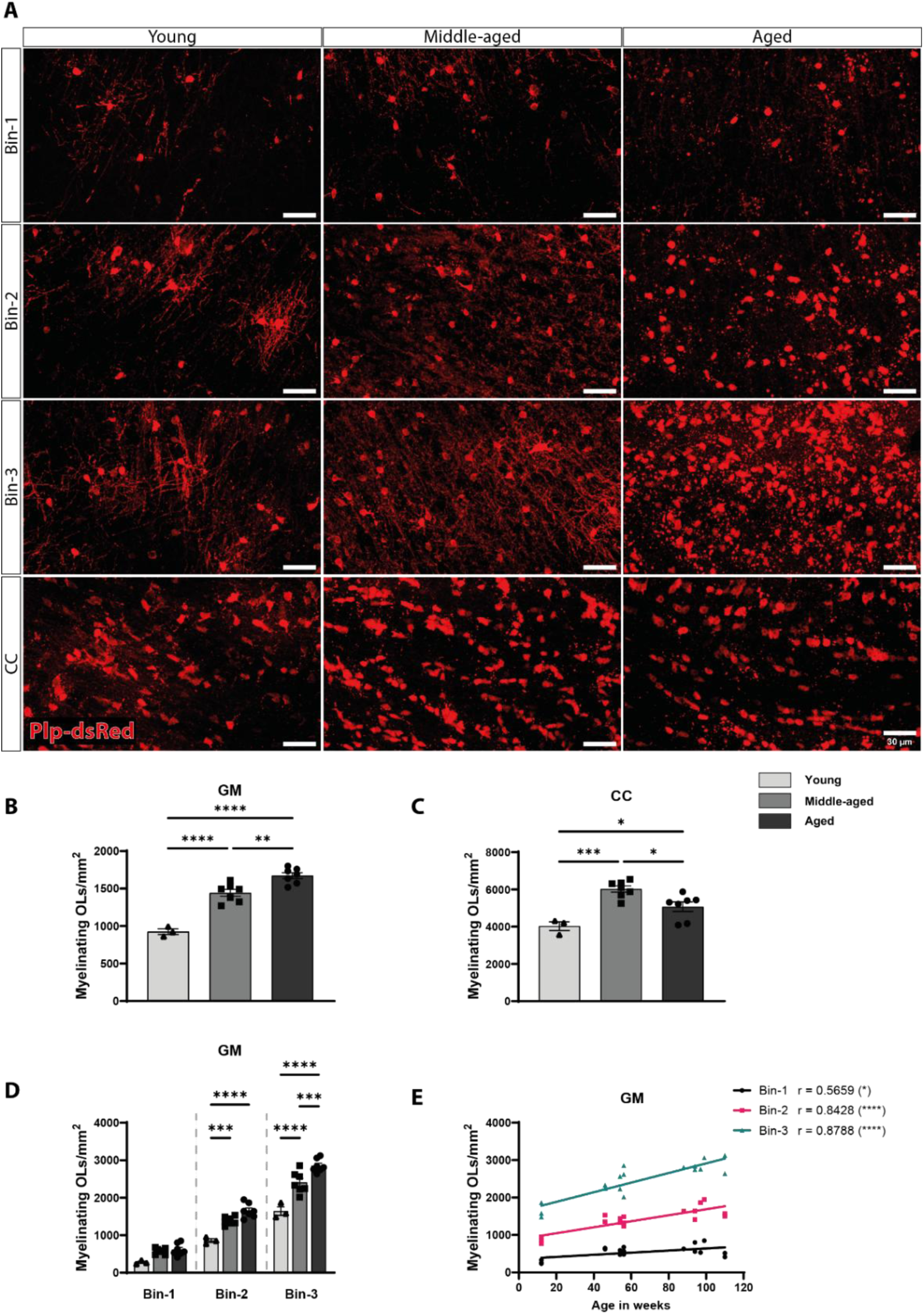
The implications of aging on myelinating oligodendrocytes are region-dependent. (A) Immunohistochemical staining of myelinating OLs (dsRed) in the reported mouse line Plp-dsRed from young, middle-aged and aged mice. The images show the different densities of myelinating OLs in bin-1 (superficial), bin-2 (intermediate) and bin-3 (deep) areas of the GM as well as the CC. The scale bar represents 30µm. (B) Numbers of myelinating OLs in the GM and (C) in the CC. (D) Numbers of myelinating OLs in different bins of the GM. (E) Simple linear regression with Pearson’s correlation analysis shows the density of myelinating OLs in different depths of the GM and their relationship to age. All data are plotted as mean ± SEM. Each dot represents an individual animal. Statistical differences were determined by unpaired one-way ANOVA with Tukey’s post-hoc multiple comparisons tests. *p* value ≤ 0.05 = *, *p* value ≤ 0.01 = **, *p* value ≤ 0.001 = *** and *p* value ≤ 0.0001 = ****.

In the CC, MOL density was also increased significantly until one year of age (young vs. middle-aged *p=0.0004*; Figure 3 C). However, in contrast to the GM, MOL numbers declined significantly in old ages (middle-aged vs. aged *p=0.0159*; Figure 4 C). This was reinforced by a non-linear lognormal fit (r^2^=0.6075; Suppl. Figure 1 E). This divergent trajectory highlights a region-specific vulnerability of mature OLs within the CC.

Layer-dependent analysis of the GM revealed that MOL accumulation occurred across the whole depth of the GM, however, to different extents. Significant increases in the numbers of MOLs were observed in bin-2 and bin-3, accompanied by a positive trend in bin-1 (young vs. aged bin-1 *p=0.0616*, bin-2 *p<0.0001*, bin-3 *p<0.0001*; Figure 4 D). Notably, MOL addition plateaued in the superficial GM (bin-1) after middle-age, while deep layers continued to accumulate MOLs into old ages (middle-aged vs. aged bin-2 *p=0.0506* and bin-3 *p=0.0002*; Figure 4 D). Pearson’s correlation analysis confirmed a significant, positive correlation between the age and the number of MOLs in all bins, with the strongest effects in bin-2 and bin-3 of the GM (bin-1: r=0.5659 *p<0.05*, bin-2: r=0.8428 *p<0.0001* and bin-3: r=0.8788 *p<0.0001*; Figure 4 E).

To determine whether apoptotic loss contributes to the age-dependent decline of MOLs in the CC but not the GM, we performed TUNEL assays. Indeed, apoptotic MOLs (TUNEL^+^dsRed^+^) were significantly more abundant in aged CC, whereas no such increase was detected in the GM (CC: young vs. aged *p<0.01*, middle-aged vs. aged *p<0.*05; Suppl. Figure 3 F-G). These findings indicate that, unlike in the GM, the CC undergoes substantial loss of MOLs with advancing age, driven at least in part by increase MOL apoptosis, underscoring the heightened susceptibility of this myelin-dense white matter tract.

### Callosal microglia exhibit an age-dependent increase in density and proliferative activity

Inflammaging is a major driver of CNS aging and is largely mediated by microglia (Mészáros et al., 2020). We therefore first assessed age-associated changes in microglial density in both regions. Interestingly, in the cortical GM, microglial numbers remained remarkably stable across all age groups, with no significant differences in their density (Figure 5 A-B). In contrast, the CC displayed a pronounced age-dependent increase in microglial density (young vs. aged *p=0.0001*, middle-aged vs. aged *p=0.0004*; Figure 5 A-C). Pearson analysis also confirmed a significant positive age-dependent correlation exclusively in the CC (CC: r=0.7891; Suppl. Figure 4 A-B). Layer-resolved analysis further showed no age-related differences in microglial density within superficial, intermediate, or deep GM bins (Suppl. Figure 4 C-D).

**Figure 5:**
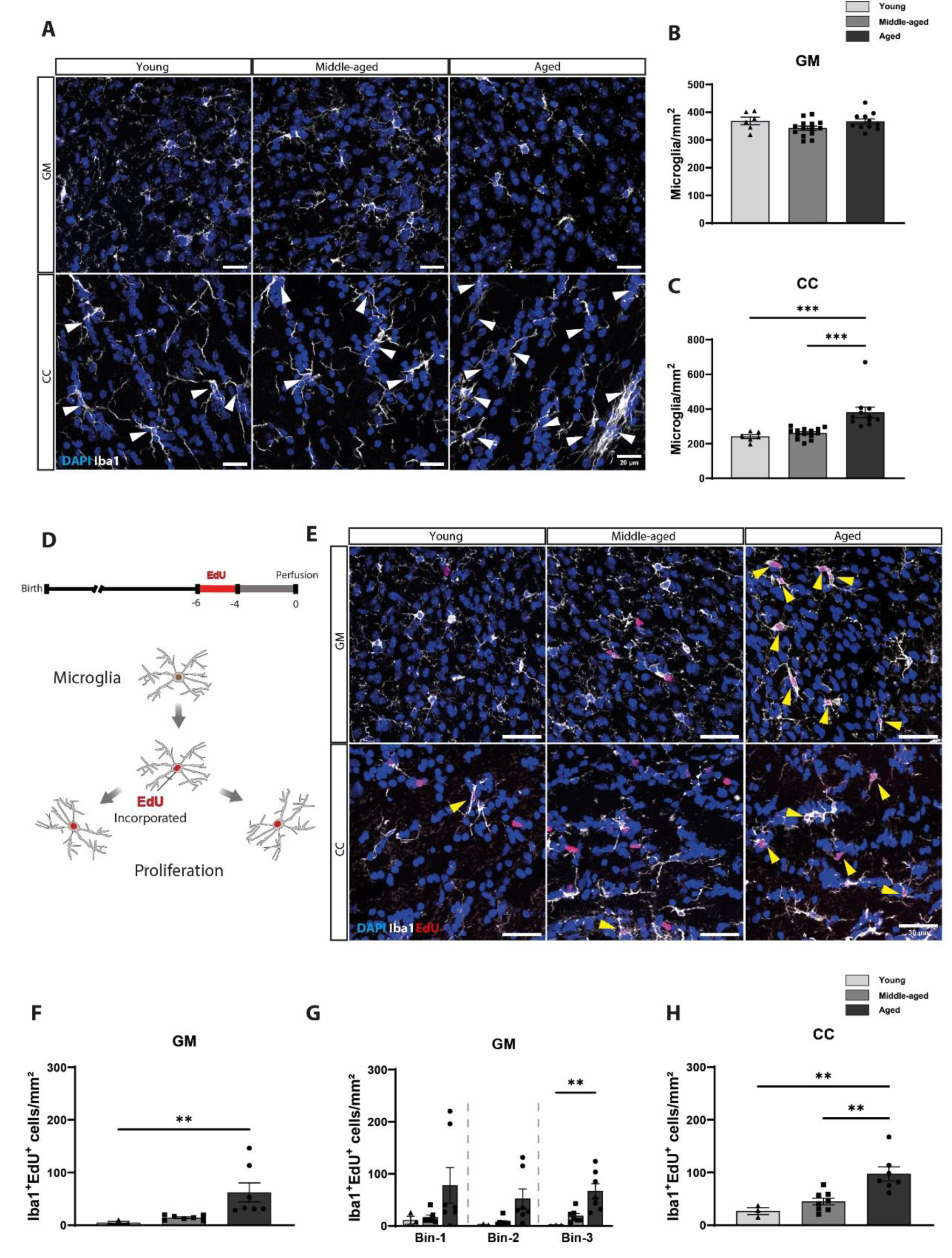
Microglia density and reactivity increase with aging in myelin-dense regions. (A) Immunohistochemical staining of microglia (Iba1^+^) and nuclei (DAPI⁺) from young, middle-aged and aged mice. White arrowheads point to microglia. The scale bar represents 30µm. (B) Numbers of microglia in the GM and (C) in the CC. (D) A schematic view of the EdU pulse (two weeks) in red and the EdU retention period (four weeks) in gray. The scheme illustrates the incorporation of EdU into the DNA of dividing microglia. (E) Immunohistochemical staining showing microglia (iba1), nuclei (DAPI), and proliferated cells (EdU) from young, middle-aged and aged mice. Yellow arrowheads point to proliferated microglia. The scale bar represents 30µm. (F) Numbers of proliferated microglia in the complete GM, (G) in different bins of the GM, and (H) In the CC. All data are plotted as mean ± SEM. Each dot represents an individual animal. Statistical differences were determined by the Kruskal-Wallis test with Dunn’s post-hoc multiple comparisons tests for graphs (F-G) and by unpaired one-way ANOVA with Tukey’s post-hoc multiple comparisons tests for graphs (B-C and H). *p*-value ≤ 0.01 = ** and *p*-value ≤ 0.001 = ***.

Given that microglial reactivity is often accompanied by altered proliferative behavior, we quantified EdU-incorporating microglia following a 2-week EdU pulse followed by 4-week retention. Interestingly, in the GM, proliferated microglia (Iba1^+^EdU^+^) increased significantly in aging animals compared with young and middle-aged groups (young vs. aged *p=0.0034*, middle-aged vs. aged *p=0.0209*; Figure 5 F). This increase was most pronounced in the deep GM (bin-3), with a strong trend in bin-2 (bin-3: young vs. aged *p=0.0031*; bin-2: young vs. aged *p=0.0569*; Figure 5 G). In the CC, proliferated microglia (Iba1^+^EdU^+^) also rose markedly with age (young vs. aged *p=0.0044*, middle-aged vs. aged *p=0.0047*; Figure 5 H).

To determine whether increasing microglial density in the CC reflected peripheral immune cells recruitment, we quantified the CD45^+^Iba1^−^ cells in the GM and the CC (Suppl. Figure 4 E). Infiltrating immune cells increased significantly with age, again restricted to the CC (Suppl. Figure 4 F-G). In summary, these findings demonstrate a pronounced age-dependent inflammatory response in myelin-rich regions, reflected by increased microglial density, heightened proliferative activity, and selective immune infiltration in the CC and deep GM layers.

### Microglia exhibit generalized age-dependent dystrophy

During our analysis of microglial density, we observed evident morphological alterations, including process retraction and the formation of cytoplasmic enlargements or “pockets” (Figure 6 A). These pockets were predominantly, but not exclusively, found near the cell body. Quantification revealed a progressive increase in the number of pocketed microglia showcasing this phenotype (young vs. middle-aged *p<0.0001*, middle-aged vs. aged *p=0.0089*, young vs. aged *p<0.0001*; Figure 6 B).

**Figure 6:**
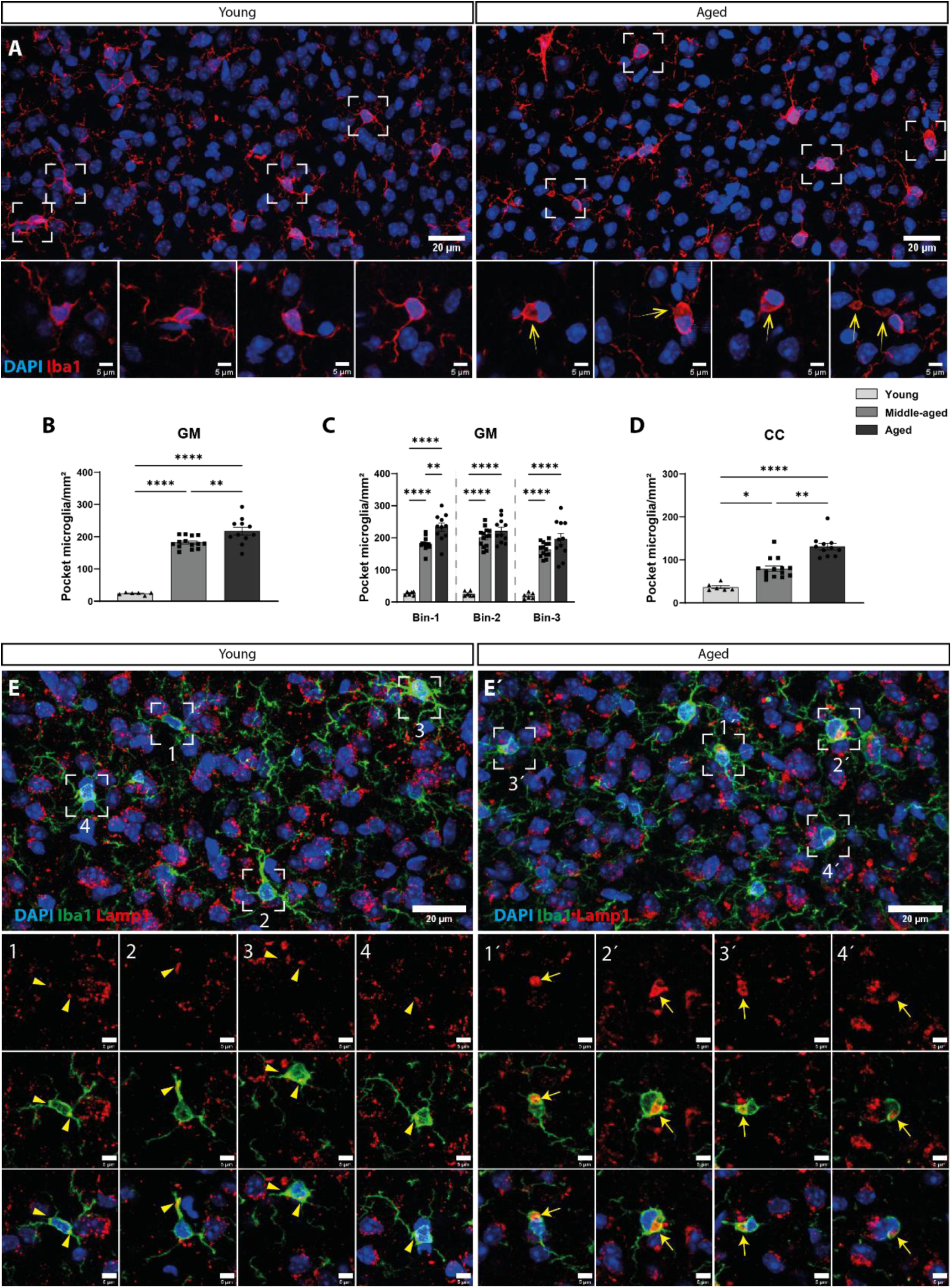
Microglia undergo generalized morphological shift and accumulation of lysosomes. (A) Representative immunofluorescence overview of microglia (Iba1⁺) and nuclei (DAPI⁺), shown as maximum intensity projection of the z-stack. Insets display zoomed-in regions of selected areas with a few individual z-planes to illustrate detailed microglial morphology. Yellow arrows point to microglia showing cytoplasmic enlargement (pocketed microglia). (B) Numbers of pocketed microglia in the GM, (C) Their distribution in different bins of the GM and (D) in the CC. (E-É) Immunohistochemical staining of microglia (Iba1^+^), lysosomal membrane (Lamp1^+^), and nuclei (DAPI^+^) from young and aged mice subsequently. Yellow arrowheads show the lysosomal distribution within microglia. Arrows show the accumulation of lysosomes within the pocketed microglia. The scale bar represents 20µm and 5µm for the overviews and insets subsequently. Data are plotted as mean ± SEM. Each dot represents an individual animal. Statistical differences were determined by unpaired one-way ANOVA with Tukey’s post-hoc multiple comparisons tests. *p* value ≤ 0.05 = *, *p* value ≤ 0.01 = **, *p* value ≤ 0.001 = *** and *p* value ≤ 0.0001 = ****.

Layer specific analysis of the GM indicates that pocketed microglia increased across all bins, peaking in middle age in bin-2 and bin-3 and plateauing thereafter, whereas superficial layers continued to rise with age (bin-1: young vs. middle-aged *p<0.0001,* young vs. aged *p<0.0001,* middle-aged vs. aged *p=0.0025*; bin-2: young vs. middle-aged *p<0.0001,* young vs. aged *p<0.0001*; bin-3: young vs. middle-aged *p<0.0001,* young vs. aged *p<0.0001*; Figure 6 C). In the CC, pocketed microglia also increased progressively across all ages (young vs. middle-aged *p=0.0493,* middle-aged vs. aged *p=0.0078,* young vs. aged *p<0.0001*; Figure 6 D). Immunostaining for the lysosomal membrane marker Lamp1 revealed that cytoplasmic pockets on aged microglia were enriched with lysosomes, contrasted with the thin-process lysosomal distribution observed in young animals (Figure 6 E and Suppl. Figure 5 G), suggesting a strong association between pocket formation and lysosomal accumulation during aging.

We next assessed myelin engulfment using the Plp-dsRed mouse line (Figure 7 A and Suppl. Figure 5 H). Engulfment events -defined as pocketed microglia containing dsRed signal- increased markedly with age in the GM (young vs. middle-aged *p=0.0002*, young vs. aged *p<0.0001* and middle-aged vs. aged *p=0.0061*; Figure 7 B). Notably, in the GM, engulfment events continued to rise beyond middle age in the deep layers (bins 2-3) (bin-2: middle-aged vs. aged *p=0.0034*, bin-3 middle-aged vs. aged *p<0.0001;* Figure 7 C), whereas pocket formation in microglia plateaued earlier (Figure 6 C).

**Figure 7:**
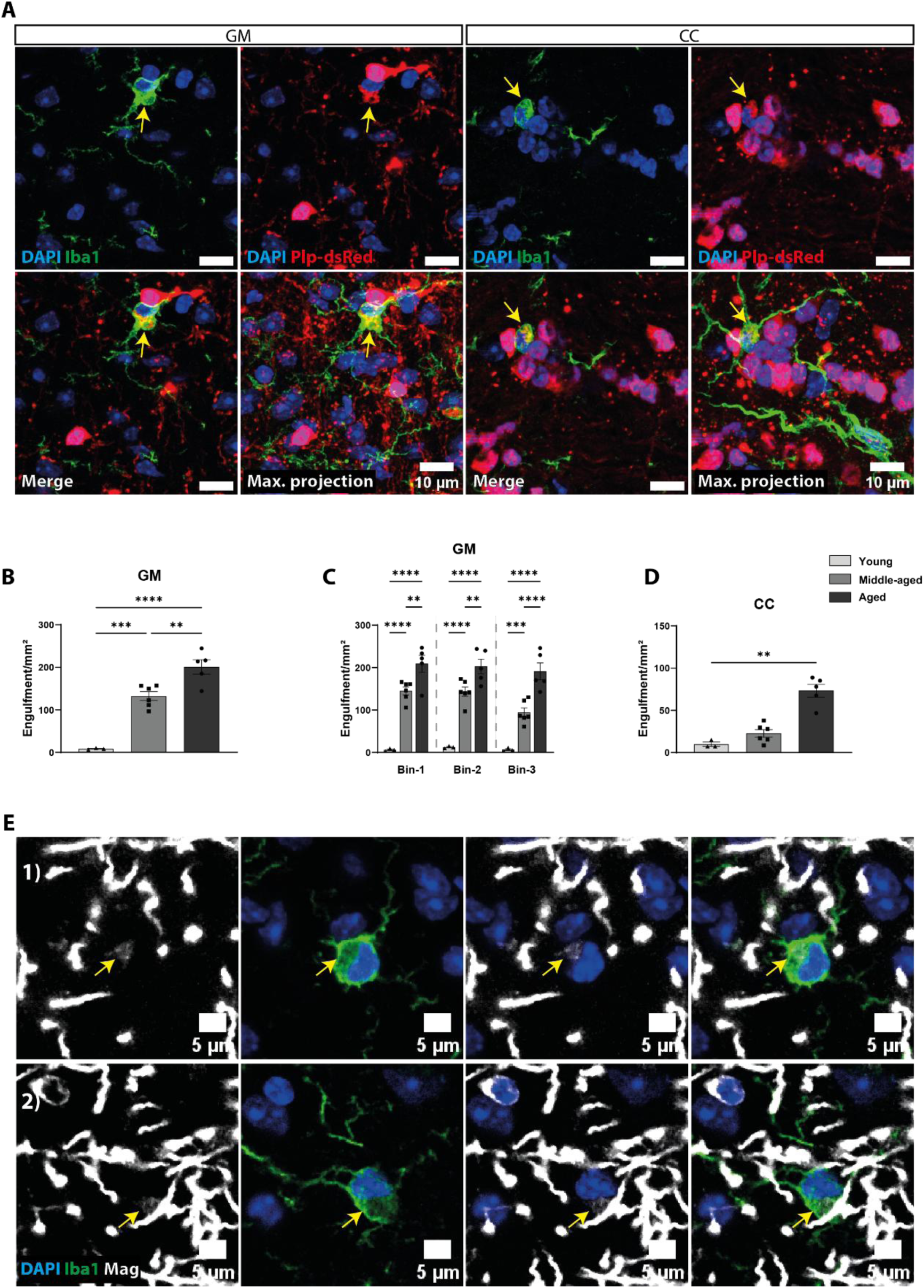
Microglia accumulate myelin debris with age. (A) Immunohistochemical staining of microglia (Iba1^+^), myelinating OLs (dsRed^+^) and nuclei (DAPI^+^) of an aged mouse from the GM and CC. Yellow arrows show Plp-dsRed reactivity within the pocket formation (engulfment). Images are shown with a few individual z-planes to illustrate detailed microglial morphology as well as maximum projection of the whole z-stack. The scale bar represents 10µm. (B) Numbers of engulfments in the GM and (C) their distribution in different bins of the GM and (D) in the CC. (E) Exemplary images from immunohistochemical staining of microglia (Iba1^+^) and myelin (Mag^+^). Yellow arrows show engulfment of myelin debris within the pockets of microglia. The scale bar represents 5µm. Data are plotted as mean ± SEM. Each dot represents an individual animal. Statistical differences were determined by unpaired one-way ANOVA with Tukey’s post-hoc multiple comparisons tests. *p* value ≤ 0.05 = *, *p* value ≤ 0.01 = **, *p* value ≤ 0.001 = *** and *p* value ≤ 0.0001 = ****.

In the CC, we likewise observed a significant increase in engulfment events in the aged group compared to the young group and a strong tendency compared to the middle-aged group (young vs. aged *p<0.0001*, middle-aged vs. aged *p<0.0001*; Figure 7 D).

To corroborate these findings, we performed an analogous analysis using the myelin marker MAG and detected clear engulfment of myelin within microglia pockets (Figure 7 E).

We then examined the proportion of microglia exhibiting pocketed morphology or engulfment events. The fraction of pocketed microglia plateaued by middle age (GM: young vs. middle-aged *p<0.0001,* young vs. aged *p<0.*0001; CC: young vs. middle-aged *p<0.05*, young vs. aged *p<0.*0001; Suppl. Figure 5 A-B). In contrast, the fraction of microglia displaying engulfment events continued to increase significantly with age in both GM and CC (GM: young vs. middle-aged *p<0.001,* young vs. aged *p<0.*0001, middle-aged vs. aged *p<0.05*; CC: young vs. middle-aged *p<0.01*, middle-aged vs. aged *p=0.053*; Suppl. Figure 5 C-D). This pattern was further supported by a non-linear lognormal curve fit, which more accurately described the relationship between age and the number of pocketed microglia compared with a linear model (r^2^=0.856 bin-1, r^2^=0.832 bin-2, and r^2^=0.758 bin-3; Figure supple. 5 E). Conversely, age and engulfment events showed a strikingly linear correlation (r=0.90 *p<0.*0001 bin-1, r=0.90 *p<0.*0001 bin-2, and r=0.94 *p<0.*0001 bin-3; Figure supple. 5 H).

Together, these data demonstrate that aging microglia undergo pronounced morphological dystrophy, which precedes functional impairment in debris clearance. While structural alterations plateau in middle age, myelin debris accumulation continues to rise, revealing a progressive impairment in microglial phagocytic capacity.

### Myelin loss is accompanied by nodal-paranodal alterations in the aging corpus callosum

Building on our observations of myelinating oligodendrocyte (MOL) loss and the pronounced inflammatory shift in the CC, we next assessed the structural integrity of myelin-associated axonal domains. Immunostaining for Caspr, a well-established paranodal marker, and the voltage-gated sodium channel Nav1.6, highly enriched at the nodes of Ranvier (Figure 8 A), revealed clear age dependent alterations.

**Figure 8.**
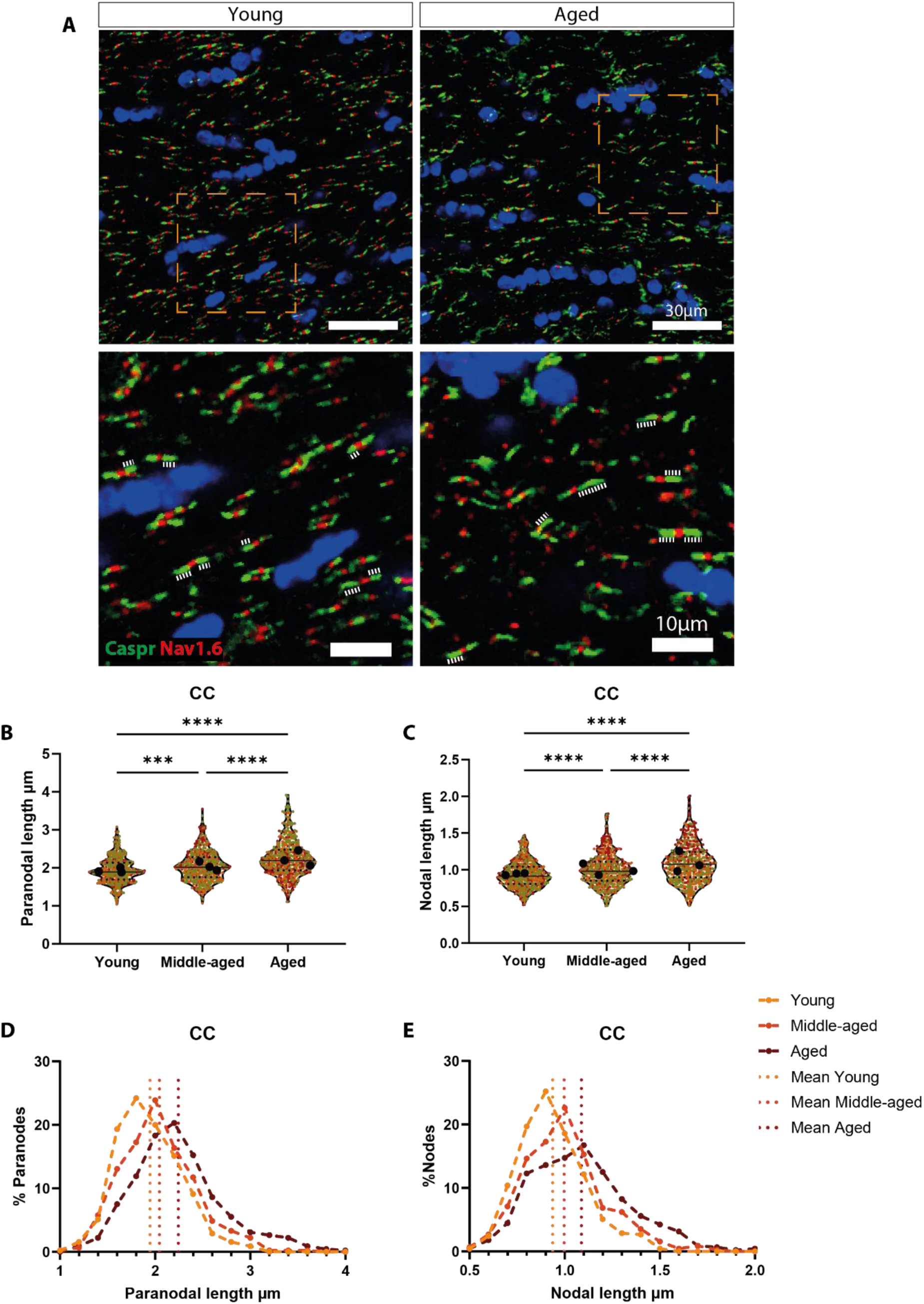
Alterations in nodal and paranodal organization with age: (A) Immunostaining for the paranodal marker contactin-associated protein (Caspr; green) and the sodium voltage channel Nav1.6 (red). Dashed lines show the Caspr^+^ paranodal domains. (B) Quantification of the paranodal length and (C) Nodal length in the CC. (D) Frequency distributions of paranodal and (E) Nodal lengths across age groups. (B-C) data are plotted as mean ± SEM. Color-coded points represent an individual node or paranode n=∼150 per mouse. Black dots indicate the average of each biological replicate n=3 for each age group.

Quantitative analysis demonstrated significant elongation of both paranodal and nodal compartments with age. Paranodal length increased progressively (young vs. middle-aged *p=0.0001*, middle-aged vs. aged *p<0.0001* and young vs. aged *p<0.0001*; Figure 8 B). Nodal length showed a similar subtle but significant increase across age groups (young vs. middle-aged adjusted *p<0.0001*, middle-aged vs. aged *p<0.0001* and young vs. aged *p<0.0001*; Figure 8 C).

Frequency distribution analysis revealed that paranodal lengths in young mice were unimodally centered on shorter lengths, whereas aging induced a progressively broader, more asymmetric distribution with increased skewness and kurtosis, indicating greater elongation and heterogeneity (Figure 8 D, skewness: 0.424, 0.577, and 0.734; kurtosis: 0.359, 0.520, and 0.835 in young, middle-aged, and aged, respectively). In contrast, nodal length distributions showed comparable skewness across age (Figure 8 E, 0.52, 0.62, and 0.583 in young, middle-aged, and aged, respectively), indicating that despite nodal elongation and variability, the relative proportion of short versus long nodes was largely preserved. These findings indicate that age-related OL loss, impaired myelin turnover, and the accompanying local inflammatory state in the CC are associated with structural remodeling of myelin–axonal domains, potentially contributing to altered axonal conduction and white matter dysfunction.

## 4. Discussion

Aging of the brain is accompanied by coordinated yet regionally heterogeneous alterations in oligodendroglial and microglial compartments, contributing to the decline in various brain functions (Wang et al., 2020). However, it remains unclear whether these age-related alterations occur independently of myelin content, a factor that varies regionally and is often overlooked in aging research. Here, we showed that age-related alterations in oligodendrocyte lineage cells and the inflammatory state of the brain strongly correlate with regional myelin content. Notably, we identified the corpus callosum (CC) and the deeper layers of the cortical gray matter (GM) as selectively vulnerable regions where we could observe NG2-glia density decline, disrupted oligodendrogenesis, increased microglial reactivity, and progressive myelin structural changes. Together, our findings reveal a tightly interconnected multicellular aging trajectory that manifests earlier and more prominently in myelin-rich regions and ends in the structural disorganization of nodal and paranodal axo–glial domains.

NG2-glia maintain homeostatic control over their density through proliferation and migration, compensating when neighboring cells die, differentiate or are experimentally ablated (Hughes et al., 2013). This homeostatic capacity supports oligodendrogenesis and myelin plasticity. However, our data revealed a loss of NG2-glia homeostasis with age, predominantly in myelin dense regions. Examining the cortical GM, which exhibits a natural myelin density gradient (Hilscher et al., 2022; Wallace et al., 2016), we observed a significant reduction of NG2-glia in the highly myelinated deep layers of the GM. Previous publications on this topic are sparse. For example, Gomez and colleagues reported a decline in the density of NG2-glia in the fimbria and the DG of the aged mouse brain (Gomez et al., 2024). In contrast, earlier studies suggested that NG2-glia density remains largely stable over time (Dimou et al., 2008b; Hughes et al., 2013; Wang et al., 2020), despite the decline of their proliferative capacity (Psachoulia et al., 2009; Wang et al., 2020). Consistent with this, our data showed a substantial decline in NG2-glia proliferation across the GM, which is insufficient to sustain their population in myelin-dense regions, leading to density loss and disrupted homeostasis. The decline in NG2-glia proliferation has been linked to cell cycle changes and alterations in metabolic and pro-mitotic pathways, which ultimately lead to the dysregulation of NG2-glia “stemness” (Leenders et al., 2024; Psachoulia et al., 2009; Rivera et al., 2021). Our findings showed that while a proliferating pool persists with age, the dynamics differ regionally. Both Ki67 and EdU analyses revealed an accelerated proliferative decline in the CC relative to the GM. This suggests that high myelin turnover and the energetic demands of dense white-matter tracts drive an early proliferative exhaustion. In the GM, although NG2-glia density decreases, the fraction of proliferating (EdU^+^) NG2-glia remained stable, indicating retained cycling capacity over extended periods. In contrast, both the number and the fraction of cumulatively and actively proliferating NG2-glia in the CC decline with age, consistent with deeper quiescence or senescence, and pointing towards a more advanced aging phenotype of NG2-glia in the CC. Indeed, single-cell RNA-seq data support this pattern, showing an increased number of quiescent NG2-glia and decreased cycling fractions with age (Heo et al., 2025), further emphasizing the emergence of senescence signatures within aging NG2-glia that could be pharmacologically targeted to rejuvenate NG2-glia by re-entering the cell cycle (Heo et al., 2025). In line with this, we could show that NG2-glia in the aged mouse express the cell cycle inhibitors CDKIs (p16, p19), which are normally found in senescent cells that undergo cell cycle halt (Safwan-Zaiter et al., 2022; Wagner et al., 2022). In support of this, a recent single-cell RNA-seq study showed a higher percentage of p16 expressing NG2-glia while aging (Gomez et al., 2024). This induction of cell cycle regulators in NG2-glia suggests that senescence-like programs increasingly constrain oligodendroglial renewal with age. This aligns with hallmarks of cellular aging that include DNA damage accumulation, metabolic stress, chromatin remodeling, and loss of niche-derived proliferative cues (López-Otín et al., 2013, 2023). The stabilization of the fraction of cycling NG2-glia in the GM, compared to the sharp decrease in the CC, indicates that regional differences in the activation of such programs may underlie divergent aging trajectories. Yet, this halt in proliferation, although not necessarily permanent, can explain the disruption of NG2-glia homeostatic control especially in areas with high demand on their differentiation and subsequently proliferation. These results reinforce emerging concepts that OPC senescence represents a major bottleneck for remyelination capacity, as demonstrated in models of chronic demyelination and in human white-matter lesions.

We could also demonstrate that aging selectively impairs the indirect mode of oligodendrogenesis, wherein NG2-glia first proliferate before they differentiate. This mechanism is critical for robust oligodendrocyte production, especially in regions of high myelin turnover, suggesting that NG2-glia lose their capacity to execute complete proliferative–differentiation sequences, consistent with a model in which cell cycle arrest precedes and impairs lineage progression. In contrast, direct differentiation remains largely preserved, suggesting that the intrinsic differentiation potential of NG2-glia is intact while the proliferative amplification is compromised. This regional vulnerability aligns with prior observations that NG2-glia in the CC proliferate and differentiate faster than those in the cortical GM (Hill et al., 2013; Viganò & Dimou, 2016b), highlighting the role of local environmental cues in shaping NG2-glia heterogeneity (Heo et al., 2025). Together, these data posit that structural myelin demand may accelerate NG2-glia aging by imposing sustained proliferative requirements that become untenable as cell cycle control deteriorates.

Oligodendrogenesis is accompanied by a continuous addition of newly formed oligodendrocytes to the myelination network (Hill et al., 2018; Rivers et al., 2008; Williamson & Lyons, 2018). Our data showed different age trajectories of mature oligodendrocytes in GM and CC, revealing an important principle of regionalized glial aging. In the GM, MOL density increases with age despite declining oligodendrogenesis, suggesting robust survival, potentially due to lower metabolic load or more favorable neurovascular and metabolic support. Conversely, in the CC, MOLs initially increase but subsequently decline in aged animals with elevated TUNEL^+^ cells, indicating a higher susceptibility to apoptosis. The high axonal density, mitochondrial activity, and myelin turnover in the CC likely amplify oxidative stress and make OLs vulnerable, further exacerbated by microglial dysfunction, implicating inflammation as a potential underlying driver. These findings explain why mammalian white matter shows disproportionate age-related atrophy compared with cortical GM (Vidal-Piñeiro et al., 2016; Wu et al., 2016).

A central finding of our study is that microglial aging is similarly influenced by the local myelin environment. In the CC, microglia exhibit increased density, proliferation, and peripheral immune cell infiltration, accompanied by morphological dystrophy, lysosomal accumulation, and impaired debris clearance – core features of microglia driven chronic low-grade inflammation, also known as inflammaging (Mészáros et al., 2020). In contrast, microglia density in the GM remains stable, but proliferation rises selectively in deeper, more myelin-rich layers, indicating that microglial activation is not driven by cortical aging per se but rather by increased burden of myelin degradation products. The inflammatory shift was more pronounced in the CC, consistent with the expansion of TREM2-dependent (Triggering Receptor Expressed on Myeloid cells 2) microglial populations reported during aging (Hart et al., 2012; Poliani et al., 2015). Microglia play a critical role in regulating myelin growth, maintenance and integrity in adulthood (McNamara et al., 2022). However, in aging, microglia become senescent, reactive and less efficient in clearing myelin debris (Thériault & Rivest, 2016). Using the Plp-dsRed mouse line, we quantified myelin inclusions within microglia cytoplasm across the lifespan and identified a striking, age-dependent increase in microglia containing cytoplasmic “pockets” characterized by lysosome-rich enlargements. Pocket formation rose steeply until middle-age and plateaued thereafter, whereas myelin engulfment continued to increase, indicating persistent phagocytic uptake despite declining degradative capacity. This expanding mismatch between engulfment and clearance mirrors metabolic overload phenotypes seen in TREM2-deficient microglia and lipid-droplet–accumulating microglia (Gouna et al., 2021; Wei et al., 2024). The accumulation of uncleared myelin debris has major functional consequences. Myelin fragments are strong inhibitors of oligodendrogenesis and myelin repair (Cunha et al., 2020), while efficient myelin clearance is required to promote remyelination (Church et al., 2017; Kotter et al., 2006; Safaiyan et al., 2016). Thus, progressive lysosomal dysfunction and debris accumulation within aging microglia create a non-permissive environment that undermines myelin maintenance, suppresses oligodendrogenesis and reinforces chronic inflammation. Together, these findings support a model in which microglial dystrophy emerges both as a consequence of elevated myelin turnover and as a driver of impaired myelin homeostasis during aging.

Myelin related changes with aging can be accompanied by disorganization of nodes of Ranvier and paranodal regions in myelinated axons (Hinman et al., 2006). In line with this, we found that loss of MOLs in the CC coincides with disorganization of axo-glial domains, including elongation and increased heterogeneity of paranodes and subtle nodal lengthening. These structural changes have significant functional consequences, as the precise structure of nodes of Ranvier and paranodal junctions is essential for saltatory conduction and action potential fidelity (Schneider et al., 2016). Similar changes were also reported under chronic neuroinflammatory conditions, where sustained microglial activation in the normal-appearing white matter of multiple sclerosis brains induced paranodal elongation, even in the absence of clear demyelination (Gallego-Delgado et al., 2020). Our data indicated that age-related MOL loss and impaired oligodendrogenesis destabilize these domains. Notably, paranodal alterations were more variable and skewed toward elongation, whereas nodal length distributions remained relatively stable. This suggests that paranodal junctions, which are structurally supported by OL processes and axo–glial adhesion complexes (Caspr/Contactin/Neurofascin-155), are particularly sensitive to OL loss and turnover deficits. These structural changes provide a mechanistic link between OL-loss, disrupted axo–glial interactions, and functional deficits in aging.

Taken together, our study reveals that aging of the brain is spatially heterogeneous, with myelin-rich regions showing accelerated NG2-glia exhaustion, impaired oligodendrogenesis, MOL loss, microglial dystrophy, and nodal/paranodal disruption. The interplay between glial aging, regional demand, and impaired myelin turnover creates a feed-forward loop that drives local white-matter vulnerability. These findings highlight the critical interplay between inflammation, myelin homeostasis, and progenitor cell function, providing a framework for understanding how cellular and structural aging converge to compromise cognitive and motor function and suggesting potential strategies to preserve white matter integrity and decelerate age-related decline.

## Acknowledgements

The authors thank Diana Giesler for her excellent technical assistance. We also thank the animal caretakers at Ulm University. Furthermore, we are grateful to Antonio Miralles Infante, Mary Wahba, Paula Klassen, Martina von der Bey, Lea Jaeger, Katrin Vollbracht, Katharina Hasenauer and Sirradu Jagana for their helpful feedback and scientific discussions.

This project was funded by the Deutsche Forschungsgemeinschaft (DFG, German Research Foundation)-Collaborative Research Center (CRC1506; ID-450627322) “Aging at Interfaces” and the Collaborative Research Center (CRC1149; ID-251293561).

## Contributions

L.D. conceived the project, designed and supervised the experiments, discussed the results, coordinated and directed the project. A.S. and L.D. designed the study. A.S. and J.Ev.B performed the *in vivo* experiments. A.S. performed the postmortem analyses, brain imaging, cellular quantifications, and statistical analyses. A.S., and L.D. wrote the manuscript.

## Supplementary Figures

**Supplementary Figure 1:**
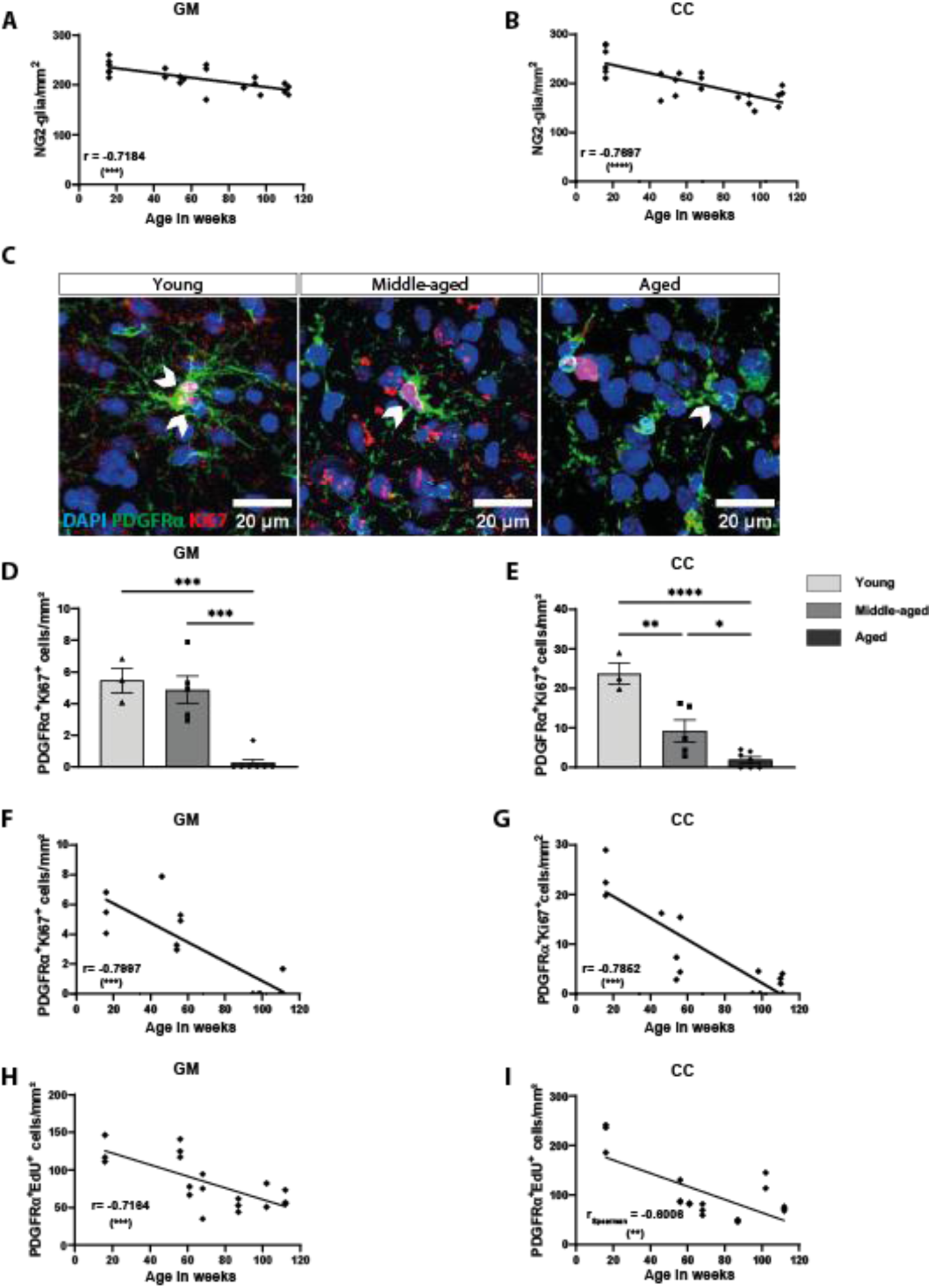
Age-dependent alterations of NG2-glia proliferation. (A) Simple linear regression with Pearson’s correlation analysis showing the density of NG2-glia with their relationship to age in the GM and (B) in the CC. (C) Immunostaining image showing active proliferation in red (Ki67^+^), NG2-glia in green (PDGFRα^+^) and nuclei in blue (DAPI^+^). White arrowheads show NG2-glia. The scale bar represents 20µm. (D) Quantifications of actively proliferating NG2-glia in the GM and (E) In the CC. (F) Simple linear regression with Pearson’s correlation analysis between age and the numbers of actively proliferating NG2-glia (PDGFRα^+^Ki67^+^) in the GM and (G) In the CC. (H) Simple linear regression with Pearson’s correlation analysis between age and cycling NG2-glia (PDGFRα+EdU+) numbers in the GM. (I) Simple linear regression with Spearman’s correlation analysis shows the density of NG2-glia and their relationship to age in the CC. Data are plotted as mean ± SEM. Each dot represents an individual animal. Unpaired one-way ANOVA determined statistical differences with Kruskal-Wallis’s post-hoc multiple comparisons tests (D-E). *p* value ≤ 0.05 = *, *p* value ≤ 0.01 = **, *p* value ≤ 0.001 = *** and *p* value ≤ 0.0001 = ****.

**Supplementary Figure 2:**
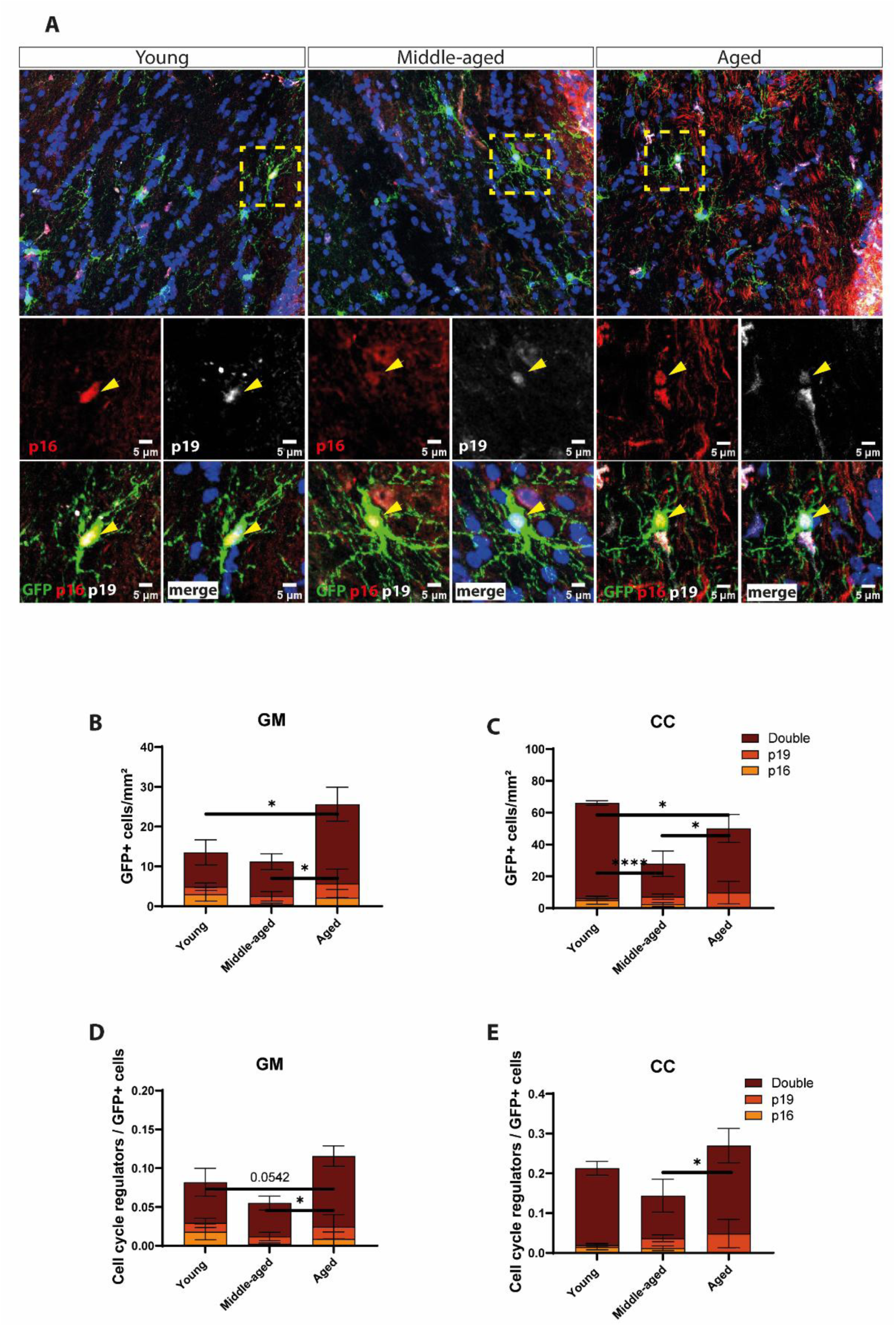
Cell cycle regulator expression in NG2-glia. (A) Immunostaining of recombined cells (GFP), cyclin-dependent kinase inhibitor (p16^INK4A+^), and p53 regulator (p19^ARF+^) from the NG2-CreERT^2^ x GFP mouse line. The dashed rectangle shows the origin of the inlets. Arrowheads indicate to GFP^+^/p16^+^p19^+^ cells. The scale bar represents 5µm. (B) The numbers of recombined cells that express the cell cycle regulators p16, p19, or both (follow color legend) in the GM and (C) In the CC. (D) The fraction of recombined cells expressing cell cycle regulator to the total recombined cells in the GM and (E) In the CC. Data are plotted as mean ± SEM. Two-way ANOVA determined statistical differences with Tukey’s post-hoc multiple comparisons tests. *p* value ≤ 0.05 = * *p* value ≤ 0.0001 = ****.

**Supplementary Figure 3:**
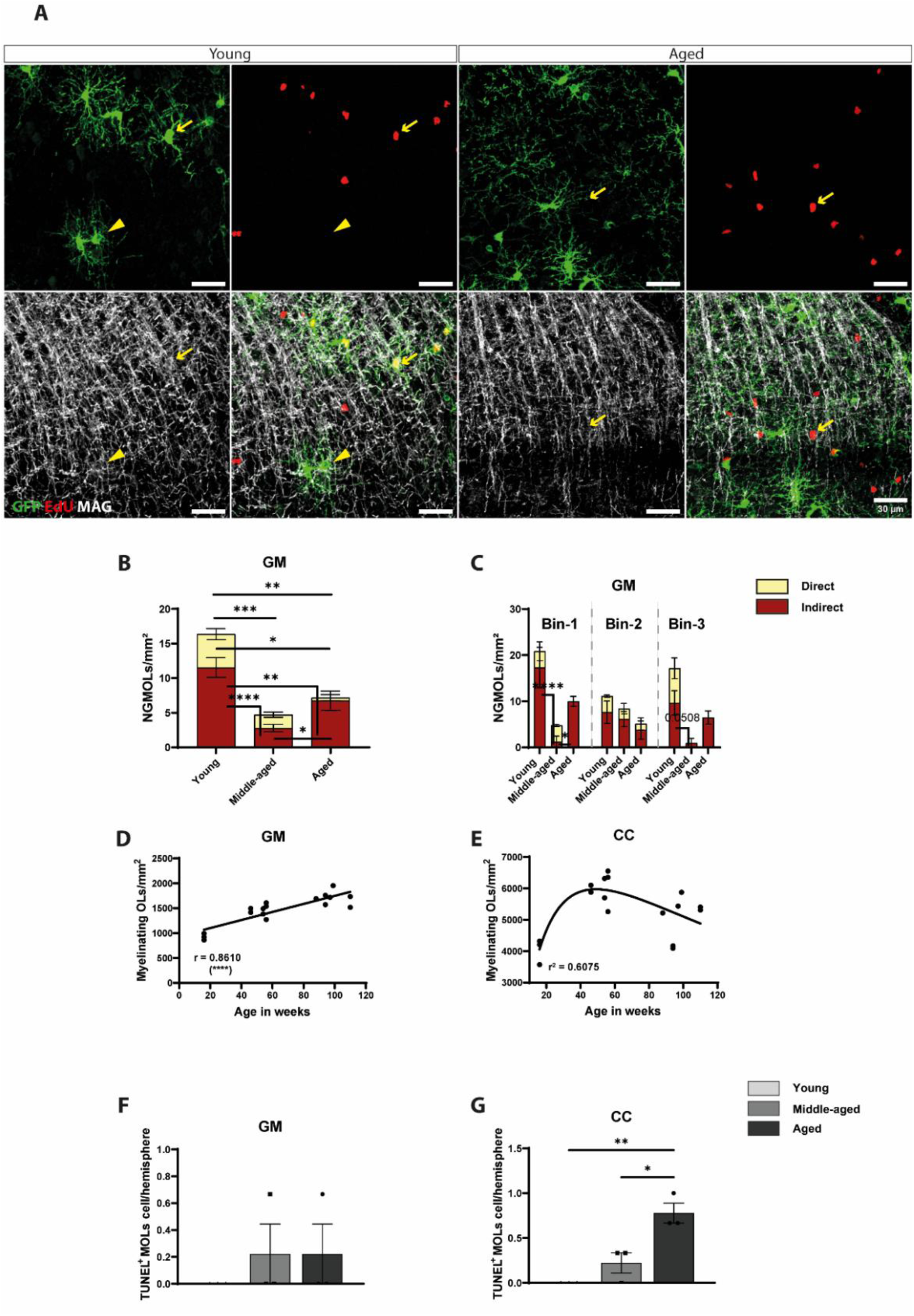
Aging effects on myelinating oligodendrocytes. (A) Immunostaining of recombined cells (GFP^+^), myelinating OLs (MOLs) (MAG^+^) and proliferated cells (EdU^+^) in the NG2-CreERT^2^ x GFP mouse line. The scale bar represents 30µm. (B) The numbers of directly (dark blue sub-column) and indirectly newly generated myelinating OLs (NGMOLs) (light blue sub-column) in the cortical GM and (C) In different bins of the GM. n=3 in young, middle-aged and aged groups. (D) Simple linear regression with correlation analysis between age and the numbers of MOLs (dsRed^+^) in the Plp-dsRed mouse line in the GM. (E) Lognormal non-linear regression fit of the numbers of MOLs across lifespan in the CC. (F) The numbers of apoptotic MOLs in the cortical GM and (G) In the CC. Data are plotted as mean ± SEM. Each dot represents an individual animal. Statistical differences were determined by two-way ANOVA (B-C) and unpaired one-way ANOVA (F-G) with Tukey’s post-hoc multiple comparisons tests. Correlation coefficient was determined by Pearson’s method (D). Goodness of the fit is shown as r² value for the non-linear lognormal fit. *p* value ≤ 0.05 = *, *p* value ≤ 0.01 = **, *p* value ≤ 0.001 = *** and *p* value ≤ 0.0001 = ****.

**Supplementary Figure 4:**
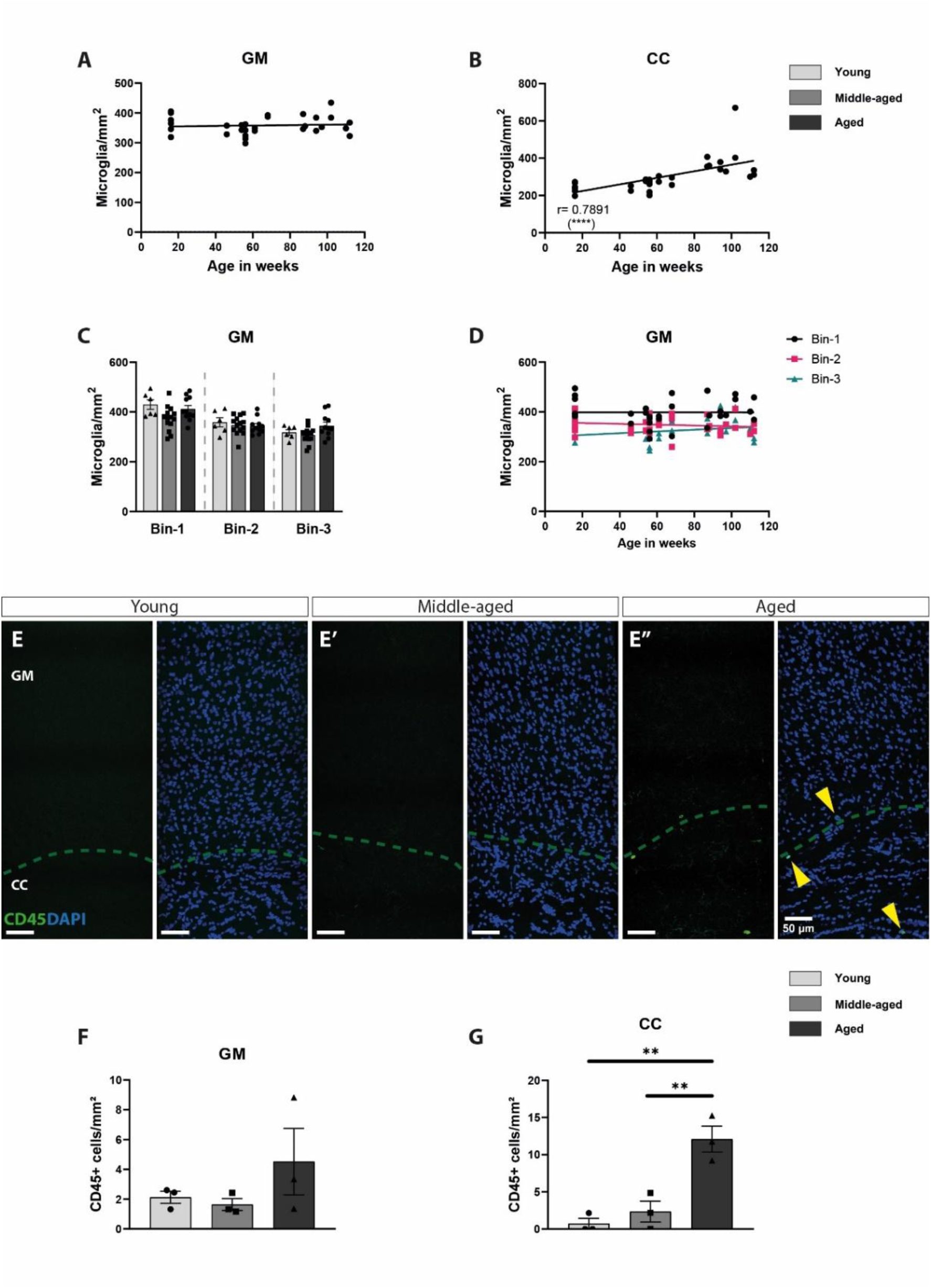
Age-dependent alterations in microglia and the blood-brain barrier integrity: (A) Simple linear regression with Pearson’s correlation analysis showing the density of microglia in relationship to age in the GM and (B) in the CC. (C) The numbers of microglia in different bins of the GM. (D) Simple linear regression showing the relationship between microglia density and age in different bins of the GM. (E) Immunostaining of leucocytes (CD45) and nuclei (DAPI). The dotted green line separates the cortical GM from its adjacent CC. The scale bar represents 50µm. (F) The numbers of CD45^+^ cells in the GM and (G) In the CC. The scale bar represents 5µm. Data are plotted as mean ± SEM. Each dot represents an individual animal. One-way ANOVA determined statistical differences with Tukey’s post-hoc multiple comparisons tests (F-G). *p* value ≤ 0.01 = ** and *p* value ≤ 0.0001 = ****.

**Supplementary Figure 5:**
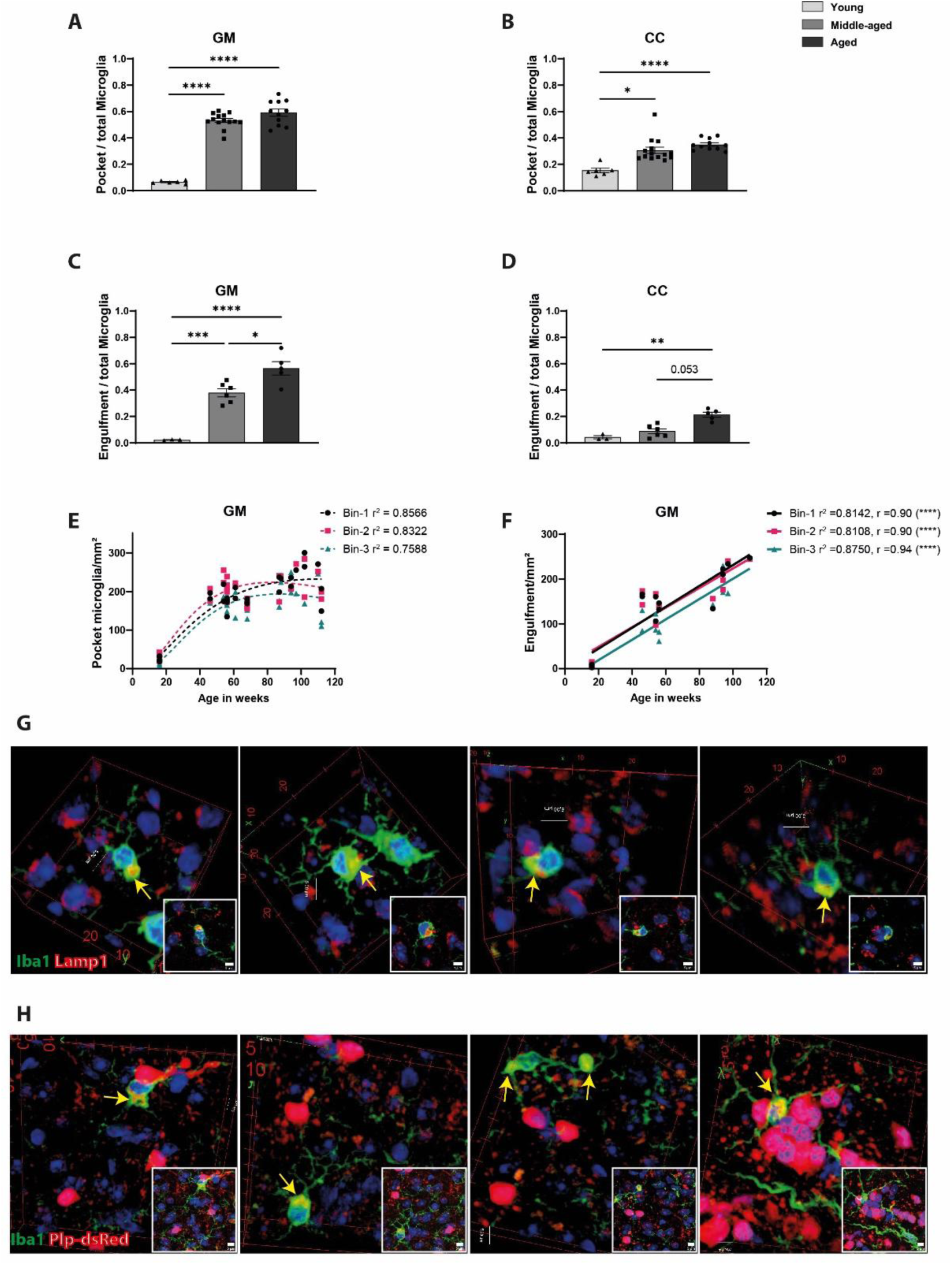
Age-dependent alterations in microglia morphology: (A) The fraction of microglia showing cytoplasmic enlargements (pockets) in the GM and (B) In the CC. (C) The fraction of microglia showing Plp-dsRed reactivity within the pockets (engulfments) in the GM and (D) In the CC. (E) A lognormal nonlinear regression model was used to fit the numbers of pocketed microglia in different bins of the GM across lifespan. (F) Simple linear regression showing the relationship between age in weeks and the numbers of engulfment events. (G) 3D projections of immunostaining of lysosomal-associated membrane protein 1 (Lamp1), microglia (Iba1) and nuclei (DAPI) from the aged mouse group. Arrows show microglia pockets filled with lysosomes. (H) 3D projections of immunostaining of myelinating oligodendrocytes and myelin (Plp-dsRed), microglia (Iba1) and nuclei (DAPI). Each snapshot is an example of what was considered as an engulfment event. Arrows show pocketed microglia containing myelin debris. The scale bar represents 5µm. Data are plotted as mean ± SEM. Each dot represents an individual animal. Two-way ANOVA determined statistical differences with Tukey’s post-hoc multiple comparisons tests (A&C). Kruskal-Wallis test determined statistical differences with Dunn’s multiple comparisons tests (B&D). Goodness of the fit was determined by r² (E). Goodness of the fit and the correlation coefficient were determined by Pearson’s r² and r subsequently (F). *p* value ≤ 0.01 = ** and *p* value ≤ 0.0001 = ****.

